# Genome-wide architecture of prolonged starvation adaptation in experimentally evolved *Drosophila* and comparative enrichment in human orthologs

**DOI:** 10.64898/2026.05.01.722137

**Authors:** Gaurav Yadav, Prachi Mishra, Ranjan Kumar Sahu, Vijendra Sharma, Pawel Michalak, Dau Dayal Aggarwal

**Affiliations:** Department of Biochemistry, University of Delhi South Campus, New Delhi, India; Department of Neurology, Houston Methodist Research Institute, Houston, Tx, USA; Department of Neuroscience, UT Southwestern Medical Center, Dallas, Tx, USA; Department of Biomedical Sciences, University of Windsor, Ontario, Canada; Institute of Evolution, University of Haifa, Haifa, Israel; Edward Via College of Osteopathic Medicine, Monroe, LA, USA; Centre for One Health Research, Virginia-Maryland College of Veterinary Medicine, Blacksburg, VA, USA

**Keywords:** Experimental evolution, starvation tolerance, genome-wide selection, polygenic adaptation

## Abstract

Long-term starvation stress represents a strong evolutionary constraint across taxa, yet the genetic architecture underlying adaptation to sustained nutrient deprivation remains poorly resolved. We experimentally evolved *Drosophila melanogaster* under starvation stress for 60 generations, maintaining four starvation-selected populations and four matched controls, and then performed whole-genome resequencing. Starvation-selected populations exhibited extended survival and pronounced genome-wide restructuring, including the expansion of low-heterozygosity regions, reduced nucleotide diversity, and recurrent sweep signatures across replicates. Drift-aware allele-frequency modeling identified 3,578 single-nucleotide polymorphisms (SNPs) whose shifts exceeded neutral expectations, indicating widespread, parallel responses to sustained starvation. Mitochondrial pathways emerged as prominent targets: nuclear-encoded mitochondrial genes were significantly enriched among low-diversity, sweep-associated, and drift-exceeding loci, and the mitochondrial origin of replication harbored a sharply differentiated variant, suggesting a mito–nuclear component of starvation adaptation. Comparative analyses further showed that human orthologs of starvation-responsive fly genes were enriched for highly differentiated variants in selected 1000 Genomes populations, with regulators of TOR/S6K signaling recurring among loci in the extreme tails of population differentiation. Together, these results define a replicate-convergent and functionally coherent genomic response to prolonged starvation in *Drosophila*, comprising regional signatures of linked selection and distributed allele-frequency shifts exceeding neutral expectations, and connect laboratory selection in flies to population differentiation in human orthologs.

## Introduction

Starvation is a pervasive ecological force that has shaped life histories, metabolic strategies, and population persistence across taxa (McCue, 2010; Rion and Kawecki, 2007), from microbes to humans (Hazan et al., 2021; Shoemaker et al., 2021). In natural populations, recurrent nutrient limitation can arise through seasonal fluctuation, famine, and ecologically marginal environments (Hoffmann and Parsons 1997; McMichael 2012), imposing sustained fitness constraints that favor molecular and physiological programs for energy conservation, storage, and stress resilience (McCue, 2010). Since starvation limits energy available for somatic maintenance, reproduction, development, and growth (Pianka, 1970; Rion and Kawecki, 2007), adaptive responses are expected to involve coordinated physiological trade-offs rather than shifts in a single trait dimension (Bronson, 1998; Shoemaker et al., 2021). Therefore, tolerance to prolonged starvation likely relies on complex genetic adaptations (Kawecki et al., 2021; Michalak et al., 2019), although the underlying genomic architecture remains largely unclear.

Natural populations of *Drosophila* harbor substantial standing genetic variation for starvation tolerance, and this trait differs significantly among populations (Aggarwal, 2014; Hoffmann and Harshman, 1999; Hoffmann and Parsons, 1997; Karan et al., 1998; Rion and Kawecki, 2007). However, its population-genetic basis remains difficult to resolve because adaptation from standing variation may leave distributed or incomplete genomic signatures, and inference in natural populations is further complicated by environmental heterogeneity and demographic history (Jain and Stephan, 2017; Kapun et al., 2020). Experimental evolution combined with whole-genome resequencing (Evolve-and-Resequence, E&R) provides a tractable alternative by imposing a defined selective regime in controlled, replicated populations, thereby enabling direct tests of genomic repeatability and allele-frequency change under selection (Garland and Rose, 2009; Kofler and Schlötterer, 2014; Schlötterer et al., 2015). E&R studies have shown that adaptive genomic architecture varies markedly across traits and selection regimes. For example, selection for accelerated development did not produce classic hard-sweep signatures or extended fixation-like intervals, consistent with incomplete responses or shifts at many loci of small effect from standing variation (Burke et al., 2010). By contrast, long-term hypoxia selection identified highly concordant fixation-like intervals concentrated in a limited number of shared genomic regions (Zhou et al., 2011), whereas a subsequent hypoxia study detected several thousand standing variants with reproducible allele-frequency shifts across replicates, while selective-sweep signatures remained rare (Jha et al., 2016). For adult starvation, however, the only prior genome-wide study reported low genome-wide heterozygosity, extended blocks of low heterozygosity, and substantial heterogeneity among selected replicates (Hardy et al., 2018). Thus, whether prolonged starvation elicits a repeatable genomic response across replicated populations, whether consistent signatures of selective sweeps emerge, and whether these signatures, alongside drift-exceeding allele-frequency shifts, define the adaptive response, remained unclear.

Long-term starvation selection also raises mechanistic questions about which functional systems are repeatedly targeted under sustained nutrient deprivation. A plausible idea is that prolonged starvation adaptation involves coordinated mito–nuclear responses rather than isolated changes in a few metabolic genes (Hill, 2015; Mossman et al., 2016; Sloan et al., 2018). Mitochondria are particularly relevant because they couple substrate utilization to ATP production, regulate redox balance, and influence stress resilience (Ballard and Melvin, 2010), all of which are central to starvation physiology. Persistent nutrient limitation may therefore be expected to affect both nuclear-encoded mitochondrial genes and the mitochondrial genome, producing coordinated differentiation across loci involved in respiration, metabolite routing, organellar maintenance, and mitochondrial genome regulation (Dowling and Wolff, 2023; Saccone et al., 2000).

The extent to which loci implicated by laboratory selection for starvation tolerance correspond to differentiated orthologs in human populations remains largely unexplored. Studies directly examining the population genetics of human starvation-related traits are absent (Benton et al., 2021; Fan et al., 2016), limiting direct comparisons. Human populations discussed in the context of recurrent famine or long-term resource limitation provide a potentially informative comparative setting, although inference remains complicated by environmental heterogeneity and demographic history (Fumagalli et al., 2011; Hancock et al., 2010; Lachance and Tishkoff, 2013). Human local adaptation often converges at the pathway level rather than at identical loci, as seen in high-altitude populations that repeatedly show differentiation in HIF-related pathways despite distinct causal variants (Alkorta-Aranburu et al., 2012; Fan et al., 2016). Within this comparative framework, asking whether starvation-responsive loci identified in experimentally evolved *Drosophila* correspond to differentiated human orthologs provides a conservative basis for evaluating broader pathway-level evolutionary correspondence.

To address these questions, we performed long-term experimental evolution under starvation stress in *D. melanogaster*, maintaining four starvation-selected populations and four matched controls for 60 generations. We measured phenotypic divergence through starvation-survival, lifespan, and lipid assays, and analyzed genome-wide variation using pooled whole-genome resequencing. We then combined complementary population-genetic analyses of heterozygosity, nucleotide diversity, selective sweep inference, and diffusion-based drift modeling to identify both regional and widespread signatures of adaptation across replicated populations. Next, we examined the functional coherence of starvation-responsive gene sets, with particular attention to mito–nuclear signatures. Finally, we tested whether human orthologs of starvation-responsive fly genes were enriched for highly differentiated variants in populations inferred to face long-term food resource limitation. This replicated approach enabled us to define the genomic architecture of prolonged starvation adaptation, distinguish selection-driven changes from neutral drift, and evaluate the repeatability and cross-species correspondence at the level of orthologous genes and pathways.

## Results

To investigate the genomic basis of prolonged starvation adaptation, we experimentally evolved *D. melanogaster* for starvation tolerance, maintaining four independent starvation-selected populations in parallel with four matched control populations (Figure 1). This replicated design provided a framework for subsequent phenotypic, physiological, and population-genomic analyses.

**Figure 1.**
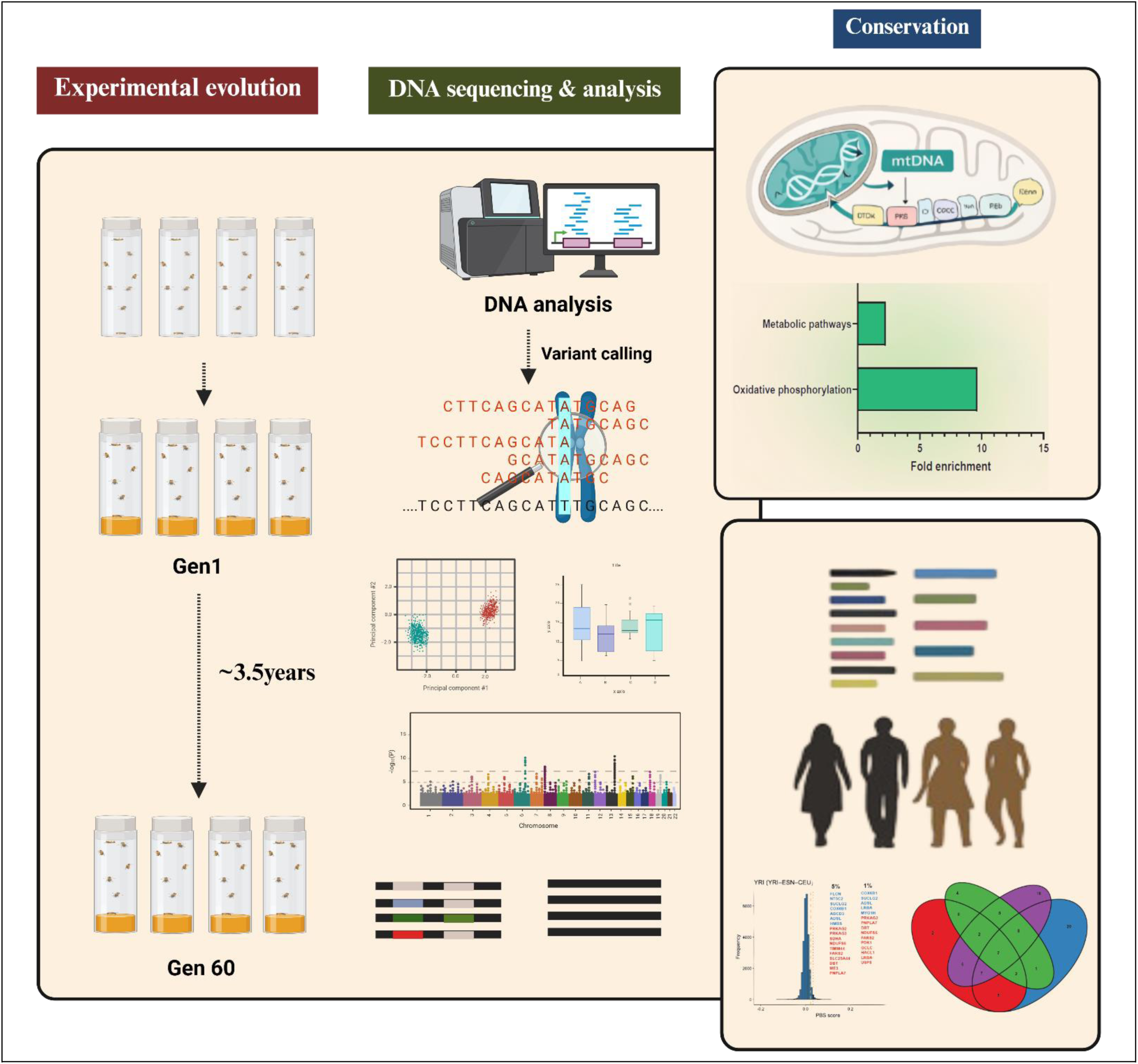
Schematic overview of experimental evolution, genome-wide DNA sequencing and analysis, and conservation-based cross-species interpretation of candidate loci.

### Genome-wide divergence under starvation selection

Starvation-selected (SS) females exhibited ∼3-fold increase in starvation survival (Nested ANOVA, *P* < 1 × 10⁻^9^) and a ∼1.5-fold increase in triacylglycerol content (*P* < 1 × 10⁻^20^) relative to controls (Figure 2A and 2B). Whole-genome resequencing of pooled females (n = 100 per population) from four starvation-selected (SS₁–₄) and four control (C₁–₄) populations, performed at high sequencing depth (>60×) and high base quality (Q > 30) (Supplementary Table S1), yielded 1,508,941 high-confidence SNPs (Supplementary Table S2). Principal component analysis (Figure 2C) revealed tight clustering of C and a clear separation of the SS populations; PC1 and PC2 explained 70.2 % of the total allelic variation (53.7% and 16.5%, respectively). Pairwise allele-frequency correlations were moderate between control and selected populations (*r* = 0.34–0.50) and stronger within SS/C replicates (*r* = 0.55–0.89) (Figure 2D). Genome-wide heterozygosity profiles (100-kb windows; 2-kb step) showed pervasive diversity loss in selected populations (mean 0.10–0.20 vs. 0.29–0.30 in C; Wilcoxon *P* = 0.02) and significantly elevated variance (SS: 0.008–0.015; C: 0.002–0.003; Levene *P* < 2.2 × 10⁻¹⁶; Figure 3A, 3B, Supplementary Figure S1, S2A and Table S3). Low-heterozygosity windows (<0.05) were abundant in SS (1,589–3,027) and rare in C populations (2–3), clustering into 29–31 contiguous blocks in SS₁–₄ populations, 11 of which were shared across all selected replicates (Supplementary Figure S2B-S2D). These shared blocks spanned ∼0.6–1.1 Mb. A similar pattern was observed for nucleotide diversity (π) (Supplementary Figure S3).

**Figure 2.**
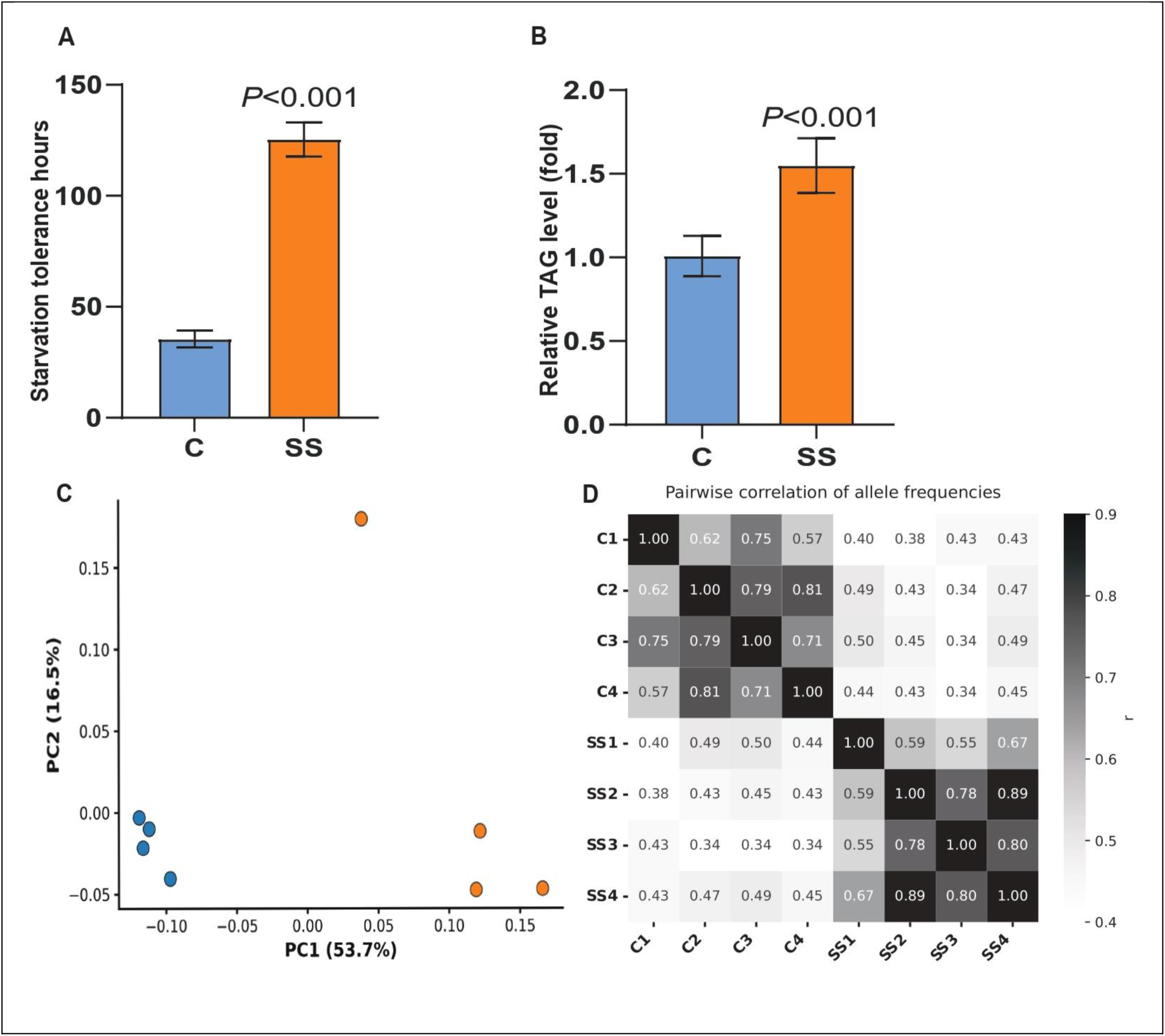
Divergence in starvation tolerance and triglyceride content, and genome-wide allele-frequency differentiation in starvation-selected versus control populations. Starvation-selected (SS) populations exhibit a pronounced extension of survival (A) under acute starvation relative to control (C) populations in females (Nested *ANOVA*, *P* < 1 × 10⁻^9^), indicating strong and repeatable phenotypic adaptation to starvation stress. (B) Relative triacylglycerol level (fold change) in SS versus C populations. Bars represent mean ± SD. (C) Principal component analysis of genome-wide allele-frequency data reveals clear and consistent separation between C and SS populations along PC1, which explains 53.7% of the total genetic variance. (D) Pairwise Pearson product–moment correlations of reference allele frequencies across all six populations. Darker shading denotes higher correlation coefficients. Control populations cluster tightly, whereas starvation-selected populations show reduced correlation with controls but high correlations among themselves.

**Figure 3.**
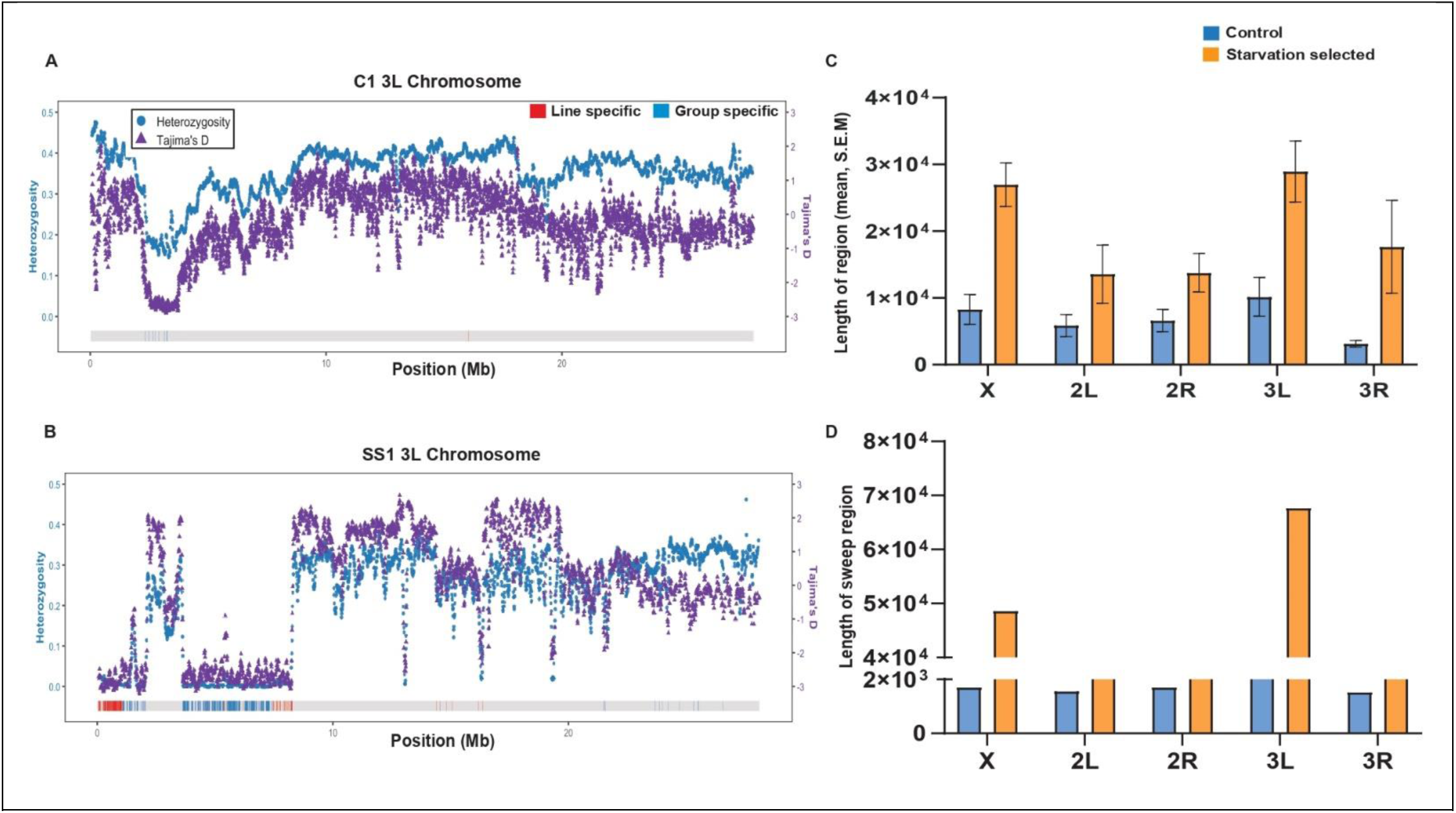
Heterozygosity and Tajima’s D plotted against selective sweep signatures along major chromosome arms of *D. melanogaster*. (A) Control (C) and (B) starvation-selected (SS) genome-wide profiles of heterozygosity (blue dots) and Tajima’s D (purple triangles) along chromosome arm 3L. Horizontal colour blocks indicate the genomic positions of inferred selective sweep regions, classified as sweep regions shared across all SS populations (blue), line-specific sweep regions detected in individual SS populations (red), and no sweep (gray). (C) Distribution of selective sweep lengths (mean ± S.E.M.) across chromosome arms (X, 2L, 2R, 3L, and 3R) in C and SS populations. (D) Lengths of shared selective sweep regions for each chromosome arm in C and SS populations

Because low-heterozygosity blocks were often extensive, they likely reflect broad consequences of linked selection rather than direct targets alone. We therefore intersected them with Pool-HMM sweep-associated regions, which capture more localized sweep structure. This combined criterion is more informative than either metric alone and enriches for intervals most consistent with directional selection within linked genomic backgrounds.

Selective-sweep inference (Pool-HMM) revealed substantially more shared sweep-associated regions in SS than in C (599 vs. 200; χ², *P* = 2.2 × 10⁻¹⁶; Supplementary Figure S4, Table S4 and Table S5), with significantly longer mean sweep lengths in SS (9.6–67.7 kb) than in C populations (1.5–2.43 kb; Wilcoxon test, *P* = 0.02; Figure 3C and 3D). Low-heterozygosity blocks reflect sustained loss of diversity, whereas sweep signals capture regions shaped by directional selection on linked variants. Their intersection identifies robust selection targets, yielding 62 high-confidence regions (hereafter referred to as sweep–low-heterozygosity regions) on chromosomes X, 3L, and 3R encompassing 255 genes, absent from controls (Supplementary Table S6 and Table S7).

Across these sweep–low-heterozygosity regions, Tajima’s D shifted from near-neutral values in controls (−0.78 to 0.75) to strongly negative values in SS populations (−3.14 to −2.37), with expected heterozygosity reduced from 0.23–0.34 to 0.001–0.055, consistent with near-complete haplotype fixation. Within these regions, some genes, for example, *lin-28*, *ND-ACP*, *mRpL50, cpo, Acat2*, and *InR*, *Sac1*, exhibited high sweep scores (>3) (Supplementary Table S8), strongly negative Tajima’s D (−2.73 to −3.14), and extreme loss of heterozygosity (0.0015–0.0028). These genes are linked to insulin/TOR signaling (*lin-28, Sac1, InR*), mitochondrial energy production (*ND-ACP, mRpL50*), and lipid β-oxidation (*Acat2*). Gowinda analysis, which corrects for gene length and SNP clustering, revealed enrichment of amino acid activation, lipid phosphatase activity, and negative regulation of insulin secretion (*P* < 0.05; Supplementary Figure S5 and Table S9). Concordant reductions in Tajima’s D and nucleotide diversity (π) across these regions provide independent support for recent, selection-driven sweeps rather than demographic effects.

### Drift-filtered genome-wide signatures

To distinguish selection-driven allele-frequency change from stochastic variation in finite populations, we applied a replicate-specific drift-filtering framework as described elsewhere (Hardy et al., 2018). This analysis identified 20,158 SNPs in SS1, 27,523 in SS2, 20,276 in SS3, and 15,407 in SS4 whose allele-frequency shifts exceeded neutral drift expectations (Supplementary Figure S6). Of these, 3,578 SNPs were shared across all four SS populations (Figure 4), indicating reproducible allele-frequency change under starvation selection. Genome-wide distributions of |ΔMAF| revealed pronounced spatial clustering of drift-inconsistent SNPs into discrete genomic regions rather than uniform dispersion, consistent with region-level selection under starvation. To further confirm whether the observed overlap among starvation-associated variants exceeded random expectation, we performed 100,000 permutation simulations by randomly sampling SNPs from the genome-wide background of 1,508,941 variants to match replicate-specific candidate sets; the observed overlap of 3578 shared SNPs lay well outside the simulated distribution (*P* = 8.29 × 10⁻^6^).

**Figure 4.**
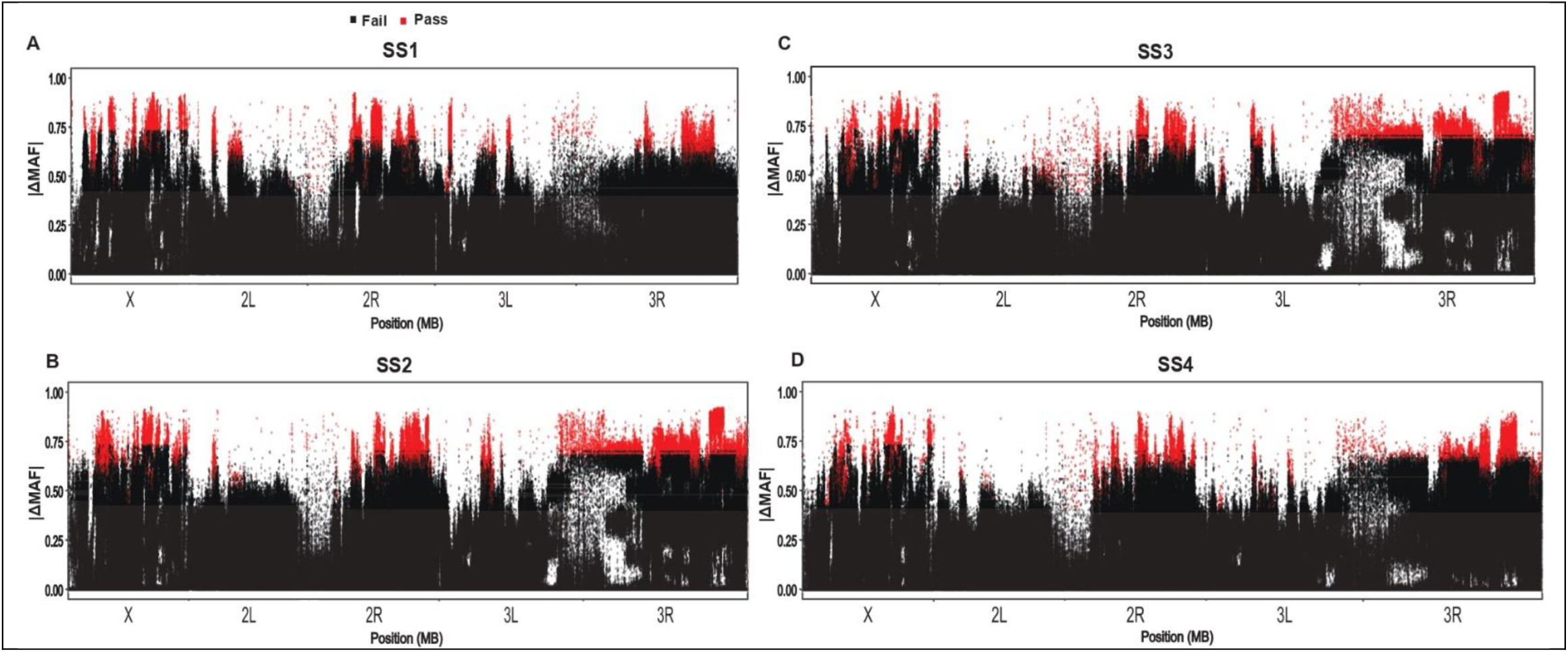
Genome-wide allele-frequency shifts after long-term starvation selection. Genome-wide minor allele-frequency changes (|ΔMAF|) are shown for each starvation-selected population (SS_1–4_) after replicate-specific drift filtering. Each point represents a single SNP, plotted by genomic position and the magnitude of allele-frequency change relative to the ancestral population. SNPs exceeding the 99.9% drift envelope are highlighted in red, whereas those within the drift envelope are shown in black. Drift-exceeding SNPs are nonrandomly distributed and cluster across multiple chromosome arms, including X, 2L, 2R, 3L, and 3R. Their recurrence in corresponding genomic intervals across independent SS replicates indicates parallel allele-frequency change under starvation selection.

Mapping the 3,578 shared SNPs identified 578 annotated genes associated with starvation-driven divergence (Supplementary Table S10). Gene Ontology enrichment analysis (using Gowinda), revealed 74 enriched categories (*P* < 0.05; Supplementary Figure S5 and Table S11), predominantly related to metabolic regulation. Enriched terms included lipid phosphatase activity and membrane lipid metabolic process, consistent with selection on lipid signaling and mobilization pathways critical for energy homeostasis during nutrient deprivation.

Only a limited number of genes (11) in the drift-filtered set overlapped sweep–low-heterozygosity regions and showed concordant population-genetic signatures, including *raskol*, *cpo*, *Ptp61F*, *par-6,* and *InR* (Supplementary Table S12). Importantly, *InR* and *couch potato* (*cpo*), both previously implicated in starvation and lipid-related traits (Schmidt et al., 2008, 2000), also showed consistent selection signatures in our dataset. All these loci exhibited elevated sweep scores (>3), shifts toward negative Tajima’s D in SS (−2.8 to −0.78 vs. −1.9 to 1.6 in C populations), and marked reductions in heterozygosity (from 0.2–0.4 in C to as low as 0.001–0.3 under SS populations). These results identify insulin/TOR signaling, growth regulation, and metabolic remodeling as major genomic features of adaptation to prolonged starvation.

### Coordinated mito-nuclear and mitochondrial signatures

Two independent nuclear gene sets (i) genes from sweep–low-heterozygosity regions and (ii) genes identified by drift-filtered allele-frequency shifts; were intersected with the *Drosophila* MitoCarta gene set to identify 33 nuclear-encoded mitochondrial genes associated with starvation-driven selection signatures (Figures 5A, 5B, and Supplementary Table S13). To test whether mitochondrial gene enrichment exceeded neutral expectations, we performed 100,000 permutation simulations using a feature-matched framework preserving chromosome and gene length. The observed overlap of 33 mitochondrial genes exceeded the null expectation of 2.45 genes, corresponding to a 13.9-fold enrichment; no permutation matched or exceeded the observed overlap (empirical *P* < 1 × 10⁻⁴). Results were robust to number-matched permutations. Many of these genes showed pronounced allele-frequency shifts between C and SS populations, with several exhibiting extreme differentiation (|ΔAF| > 0.60). Strongly differentiated loci included *PheRS-m*, *ValRS*, *mRpS30*, *mRpL50*, and *mRRF2*, which encode core components of mitochondrial translation and ribosomal function, consistent with modulation of mitochondrial protein synthesis under nutrient limitation. Similar signals were detected in genes involved in central carbon metabolism and respiratory function, including TCA-cycle and pyruvate-utilization genes (*SdhA*, *Idh3g*, *Men-b*, *Dbct*, *Pdk*), multiple components of the electron transport chain and ATP synthesis (*ND-13A*, *ND-ACP*, *UQCR-14*, *COX6B*, *ATPsynD*), and the uncoupling protein *Ucp4A*. Additional differentiation was observed in genes involved in lipid utilization and mitochondrial structure (*Mfe2*, *Abcd3*, *CLS*, *Phb2*), as well as regulators of mitochondrial dynamics and redox homeostasis (*Nipsnap*, *Miro*, *ScsbetaG*, *Alr*), indicating coordinated changes across mitochondrial metabolic and structural pathways.

**Figure 5.**
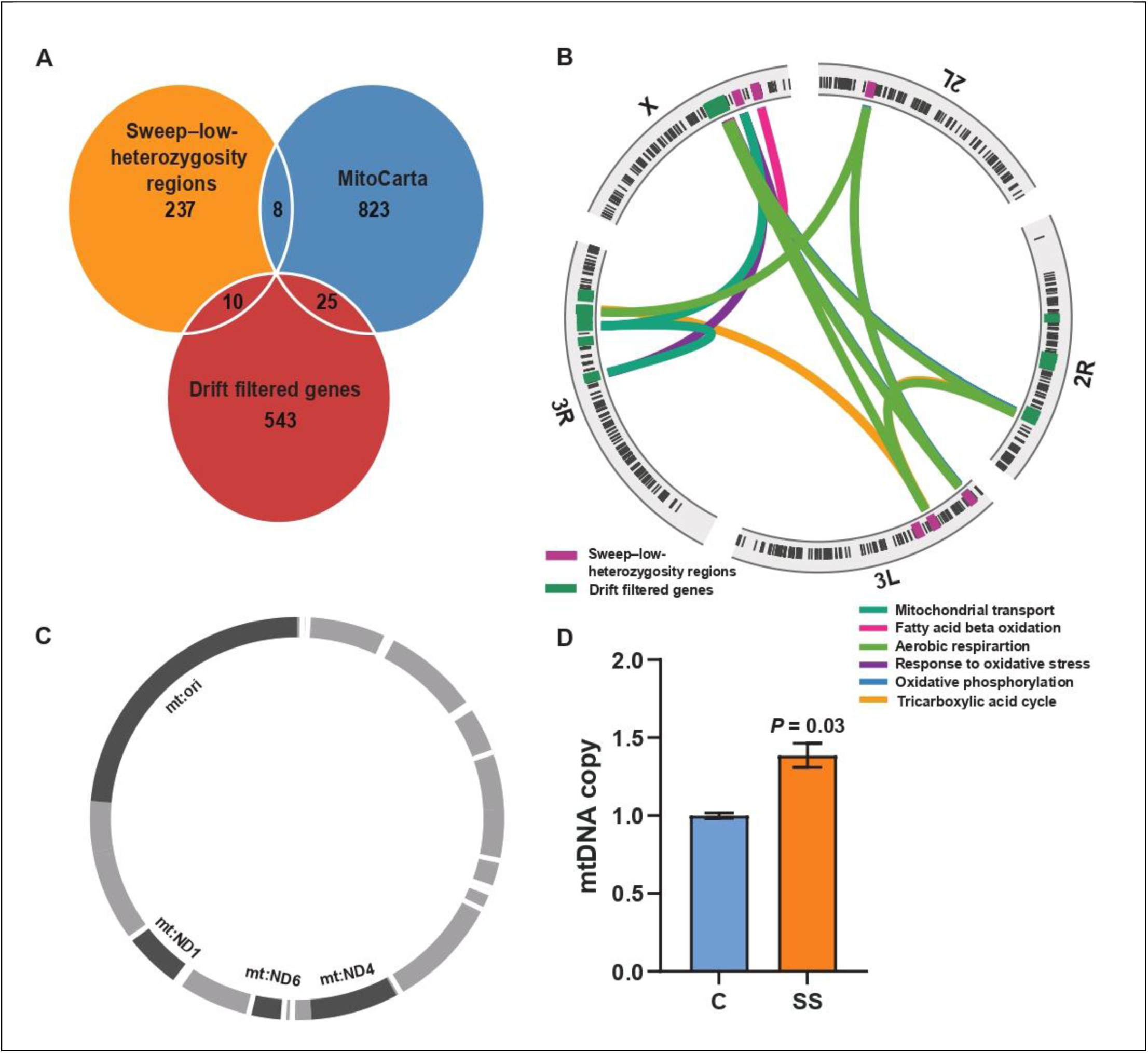
Convergent mito–nuclear and mitochondrial signatures of starvation selection. (A) Overlap among genes in sweep–low-heterozygosity regions, drift-filtered candidates, and the MitoCarta gene set. (B) Circos plot of MitoCarta genes overlapping sweep–low-heterozygosity regions (pink) or drift-filtered candidates (green); nonoverlapping genes are shown in black. Inner ribbons denote associated biological processes. (C) Mitochondrial genes harboring differentiated SNPs are highlighted in dark gray; the strongest differentiation occurs near mtORI (ΔAF up to ∼0.4). (D) mtDNA copy number is higher in starvation-selected populations than in controls (mean ± SEM; Student’s t test, *P* = 0.03).

Analysis of mitochondrial allele-frequency shifts revealed 70 segregating SNPs across the mitochondrial genome. Variants at the mitochondrial origin of replication (mtORI) showed elevated ΔAF across SS populations (Figure 5C). RNA secondary-structure modelling revealed that mtORI sequences from SS occupy a more constrained folding landscape (mean MFE = −27.12 kcal/mol) relative to C populations (−26.56 kcal/mol) (Supplementary Figure S7). While multiple mtORI regions are transcribed and also form primers for DNA replication in humans (Wanrooij et al., 2010), comparable evidence is lacking in *Drosophila.* Further, mtDNA copy-number estimates based on *cyt-b* showed a modest but reproducible increase in SS populations (*P* = 0.03; Figure 5D and Supplementary Table S14), which plausibly enhances mitochondrial function (Matsushima et al., 2004). These results support coordinated modulation of mitochondrial genome properties and nuclear-encoded mitochondrial function during prolonged starvation adaptation.

Consistent with these mitochondrial and metabolic signatures, starvation-selected populations also exhibited a significant increase in adult longevity (∼1.2-fold; Figure 6), suggesting that prolonged starvation adaptation extends beyond acute survival to broader life-history effects.

**Figure 6.**
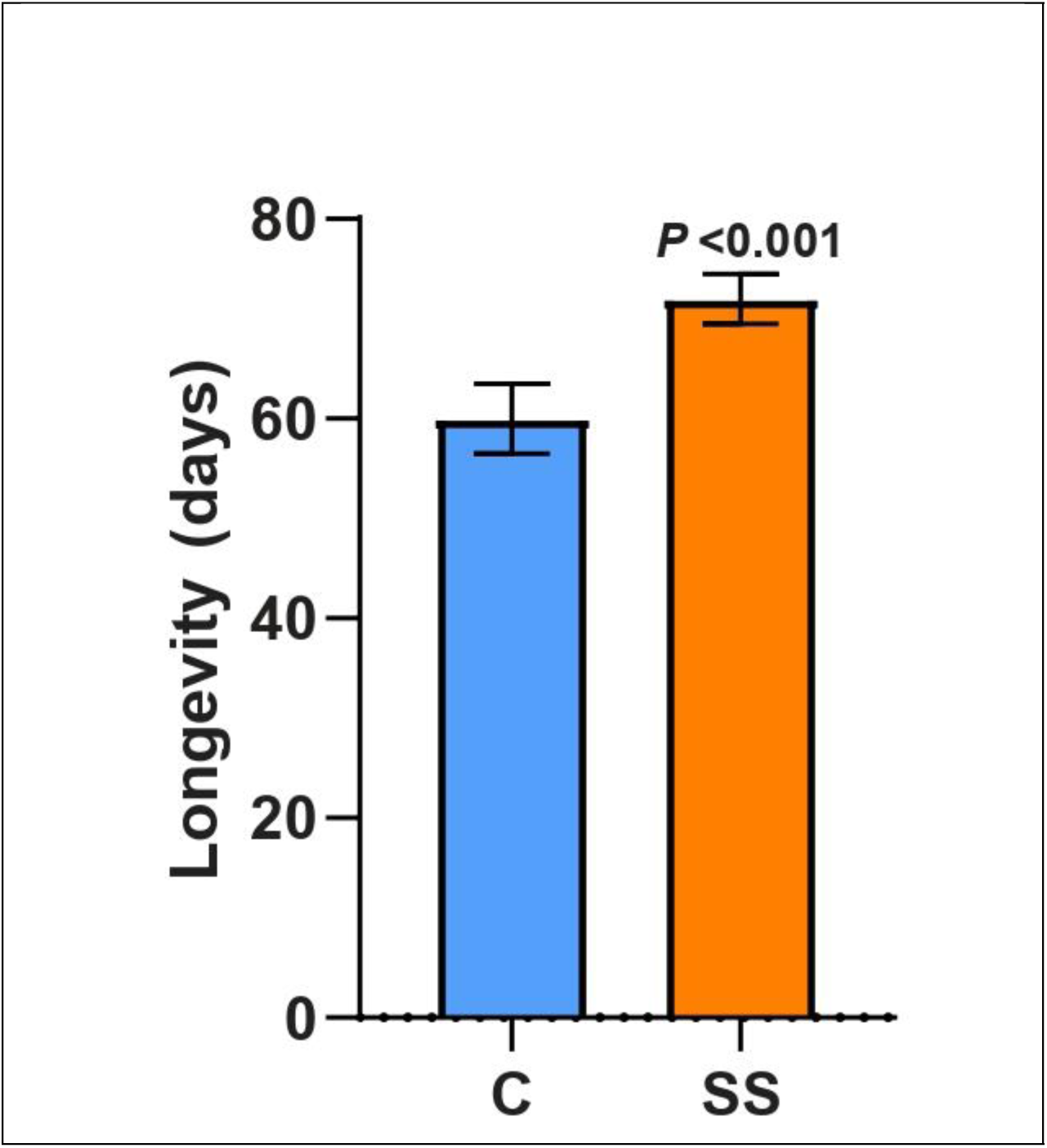
Increased longevity in starvation-selected populations. Mean adult lifespan (days) of control (C) and starvation-selected (SS) flies. Bars show means across replicate populations; error bars indicate ±SD. Starvation-selected flies lived significantly longer than controls (Nested ANOVA, *P* < 0.001).

### Comparative enrichment in human orthologs of starvation-responsive fly genes

To assess cross-species correspondence, we asked whether genes under selection in starvation-evolved *Drosophil*a also show elevated differentiation in selected human populations from the 1000 Genomes Project (see Methods). Fly genome analyses revealed two complementary classes of starvation-responsive genomic signal: 62 sweep–low-heterozygosity overlap regions encompassing 255 genes, and 3,578 drift-filtered SNPs mapping to 578 genes. Mapping these to human orthologs yielded 144 and 413 genes, respectively. We analyzed genome-wide SNP data from four focal populations: Bengali from Bangladesh (BEB), Luhya in Webuye, Kenya (LWK), Sri Lankan Tamil in the UK (STU), and Yoruba in Ibadan, Nigeria (YRI) and estimated lineage-specific differentiation using the Population Branch Statistic (PBS) within focal–reference–outgroup triplets specified in Methods (Figure 7).

**Figure 7.**
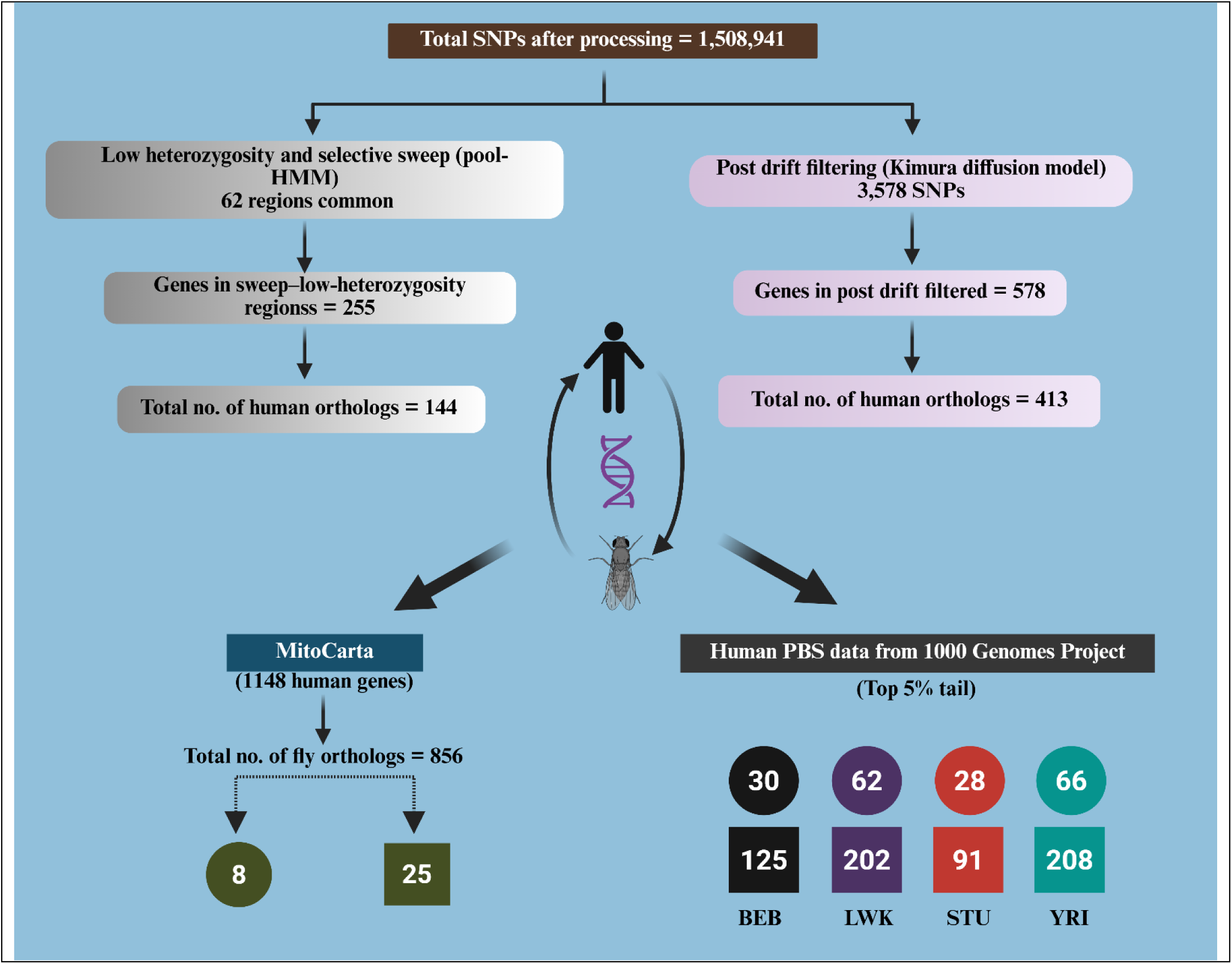
Workflow summarizing SNP filtering, ortholog mapping, and cross-species comparison with MitoCarta and human PBS candidates.

Orthologs of drift-filtered fly genes showed strong and highly significant enrichment for SNPs in the extreme tails of the PBS distribution across all four populations (Figure 8A-8D; top 5% tail: *P* = 5.54 × 10⁻⁵³, 1.81 × 10⁻⁶³, <2.5 × 10⁻¹¹⁴, and 0.0157; top 1% tail: *P* = 3.49 × 10⁻³⁵, 1.85 × 10⁻⁷, <4.71 × 10⁻¹²⁸, and 3.35 × 10⁻⁴ for BEB, LWK, STU, and YRI, respectively; binomial tests). Likewise, orthologs derived from sweep–low-heterozygosity regions showed modest but consistent enrichment, with 30–66 genes per population harbouring high-PBS SNPs in the top 5% tail, and 7 genes shared across all 4 populations (Figure 8E). Drift-filtered orthologs exhibited broader signals: 91–208 genes per population contained high-PBS variants, and 38 genes were shared across all populations (Figure 8F). Across both ortholog sets, 43 human genes were shared; notably, two genes (*raskol* and *rg*) were consistently recovered across sweep–low-heterozygosity regions and drift-filtered analyses and were also differentiated across all human populations examined (Supplementary Table S15). Notably, genes *S6k* and *sws* in *Drosophila* (*RPS6KA2* and *PNPLA6* in humans) showed concordant signals of differentiation in both the species.

**Figure 8.**
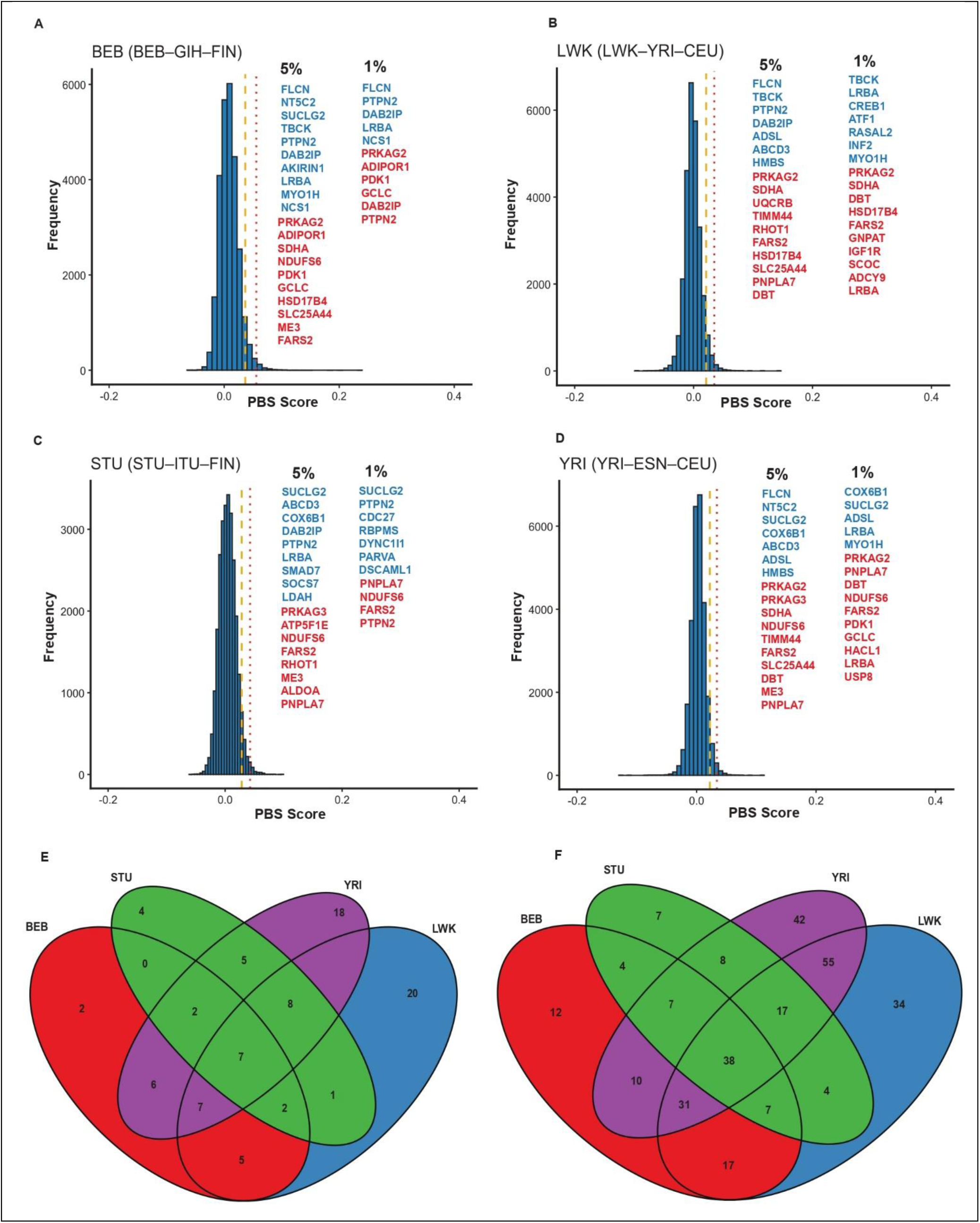
Allele-frequency differentiation in human orthologs of fly starvation-selected genes. Genome-wide PBS distributions are shown for each focal population. (A) Bengali in Bangladesh (BEB) vs Gujrati in Houston (GIH) vs Finnish in Finland (FIN), (B) Luhya in Webuye, Kenya (LWK) vs Yoruba in Ibadan, Nigeria (YRI) vs Utah residents with northern and western European ancestry (CEU). (C) Sri Lankan Tamil in the UK (STU) vs Indian Telugu in the UK (ITU) vs FIN and (D) YRI vs Esan in Nigeria (ESN) vs CEU, with vertical dashed lines indicating the 95th (top 5%) and 99th (top 1%) percentile thresholds. Human orthologs of *Drosophila* starvation-selected genes are highlighted, and many falls within the extreme upper tails of the PBS distributions. In the top 5% PBS tail, 30, 62, 28, and 66 genes overlapped with sweep–low-heterozygosity regions, and 125, 202, 91, and 208 genes overlapped with drift-filtered genes in BEB, LWK, STU, and YRI, respectively (binomial test; *P* = 5.54 × 10⁻⁵³, 1.81 × 10⁻⁶³, <2.5 × 10⁻¹¹⁴, and 0.0157 for BEB, LWK, STU, and YRI, respectively), blue indicates genes in sweep–low-heterozygosity regions; red indicates drift-filtered genes. Notably, genes involved in nutrient sensing, mitochondrial function, and lipid or amino-acid metabolism, including PTPN2, DAB2IP, PRKAG2, SDHA, FARS2, PNPLA6, DBT, and HSD17B4, exceed the 95th and 99th percentile PBS thresholds, indicating unusually strong lineagespecific differentiation at loci functionally linked to metabolic regulation. **(E)** Overlap among genes falling within the top 5% PBS tail across populations for overlapping sweep-low-heterozygosity regions. **(F)** Overlap among genes falling within the top 1% PBS tail across populations for driftfiltered genes.

As reported for hypoxia adaptation in high-altitude human populations (Alkorta-Aranburu et al., 2012; Jha et al., 2016; Zhou et al., 2013), overlap at the level of individual genes was limited, whereas enrichment at the pathway and functionally related gene set levels was significant. This pattern is consistent with polygenic adaptation, in which shared selective pressures drive coordinated shifts in allele frequencies across conserved metabolic networks rather than the repeated fixation of identical variants. Altogether, these results demonstrate that genes repeatedly targeted by starvation selection in *Drosophila* are nonrandomly enriched among highly differentiated loci in human populations experienced resource limitations, supporting conserved polygenic adaptation of metabolic pathways across species.

## Discussion

Our experimental evolution regime produced a strong starvation-adaptive phenotype, with ∼3-fold greater survival and ∼1.5-fold higher lipid storage, providing a clear physiological basis for interpreting the genomic response to prolonged starvation stress. Among 1,508,941 high-confidence SNPs, we identified two complementary classes of genomic signals: 62 regions with sweep–low-heterozygosity, and 3,578 shared SNPs with allele-frequency shifts exceeding neutral expectations after replicate-specific drift filtering. These signals mapped to 255 and 578 genes, respectively, indicating that prolonged starvation led to both localized genomic restructuring and broader, distributed changes in allele frequency. These genes were enriched for metabolic and regulatory functions, including insulin/TOR signaling, mitochondrial bioenergetics, lipid remodeling, and autophagy. Comparative analyses further revealed enrichment of highly differentiated variants in human orthologs of these genes, consistent with pathway-level similarities under putative nutritional constraints.

### Regional and distributed signatures of selection

From an evolutionary perspective, long-term exposure to starvation across generations causes persistent directional selection, with fitness differences continually expressed over time (Bubliy and Loeschcke, 2005; Harshman and Schmid, 1998; Schwasinger-schmidt et al., 2012). Under these conditions, adaptive responses are likely to heavily rely on existing genetic variation spread across multiple functionally related loci (Boyle et al., 2017; Hermisson and Pennings, 2005). Yet substantial genetic redundancy within these networks can generate heterogeneous genomic trajectories (Barghi et al., 2019); whereas variation in base-population genetic composition and starvation intensity may also favor more parallel responses at a subset of loci (Schlötterer, 2023). Strong selection should also promote pervasive hitchhiking, causing linked genomic segments to rise with beneficial alleles and thereby generating regional signatures of linked selection (Cutter and Payseur, 2013; Schlötterer et al., 2015). Consistent with this, our starvation-selected populations exhibited extensive and repeatable heterozygosity loss across large chromosomal regions, accompanied by reductions in nucleotide diversity and strongly negative Tajima’s D, indicating pronounced distortions in the site-frequency spectrum. Similar genome-wide decreases in diversity have been observed in our experimentally evolved desiccation-selected populations (Kang et al., 2016) and hypoxia-adapted *D. melanogaster* (Zhou et al., 2011). Altogether, the scale, continuity, and repeatability of these signals are more compatible with linked directional selection due to prolonged starvation than with random divergence driven solely by genetic drift.

This interpretation is further supported by diffusion-based allele-frequency modeling, which identified numerous variants with frequency shifts surpassing neutral expectations across starvation-selected populations. A previous starvation-selection study reported significant divergence among selected replicates but was limited by pronounced heterogeneity among them (Hardy et al., 2018). In contrast, the repeated appearance of both regional diversity troughs and drift-exceeding SNPs across independent replicates in our data indicates a more consistent genomic response to prolonged starvation. The limited overlap between sweep-associated diversity troughs and high-confidence drift-filtered SNPs further suggests that regional linked-selection signals and distributed allele-frequency shifts reflect partly separate components of the adaptive response. These patterns support a mixed adaptive architecture, in which strong linked selection is concentrated within localized genomic regions, while a broader polygenic component accumulates through repeatable changes in allele frequency across multiple functionally related loci.

An inherent limitation of inference under pervasive linked selection is the trade-off between the strength of the adaptive signal and fine-scale resolution (Kaplan et al., 1989; Smith and Haigh, 1974). In Pool-seq data, extensive hitchhiking obscures causal variants within the surrounding haplotypic background, as linked alleles coalesce into extended genomic blocks (Franssen et al., 2015; Schlötterer et al., 2015). Our findings on linked selection, therefore, support interpreting candidate intervals and genes as signal-bearing genomic units rather than discrete causal sites. Nonetheless, their consistent functional enrichment and recurrence across independent replicates indicate that the underlying signal remains biologically meaningful. Thus, hitchhiking does not merely limit the mapping of candidate genes; it actively influences the adaptive response by driving correlated evolution across linked loci (Frankham, 1996; Lande, 1982).

### Comparative analysis with previous starvation studies

We compared our candidate genes with those reported in three key studies: a long-term analysis of starvation-selected populations (Hardy et al., 2018), a short-term starvation-selection study conducted in the context of experimental adaptive radiation (Michalak et al., 2019), and an investigation into chronic larval malnutrition (Erkosar et al., 2023; Kawecki et al., 2021). Across these datasets, gene-level concordance was modest and highly dependent on the analytical framework used. Overlap was consistently lower for the sweep–low-heterozygosity candidate gene set (identifying only 7, 1, and 3 genes in common with ref. [21,7,8], respectively) than for the broader drift-filtered set (16, 16, and 21 genes, respectively; Supplementary Table S16). This limited overlap persisted even though both our study and Hardy et al. (Hardy et al., 2018), employed high-intensity, long-term starvation selection. These differences suggest that even under broadly similar selective regimes, populations with different genetic architectures can achieve similar phenotypic outcomes through different genomic pathways. Such a pattern aligns with polygenic adaptation under genetic redundancy (Barghi et al., 2020, 2019). In this framework, standing variation in the founding population may influence which nodes within a redundant network show the strongest genomic response, resulting in idiosyncratic regional signatures despite broadly shared phenotypic outcomes.

Part of this disparity might also reflect differences in nutritional stress levels, which can influence the dominant mode of selection. While chronic malnutrition from a nutrient-poor diet has been linked to signs of balancing selection (Kawecki et al., 2021), our findings mainly show patterns consistent with strong directional selection. A plausible interpretation is that complete starvation acts as a more rigorous selective filter than a poor diet, thus restricting the maintenance of balanced polymorphisms under milder nutritional stress. Consequently, key regulatory nodes like *InR* and *cpo* may serve as critical loci repeatedly involved in starvation adaptation. Both genes were also identified by Hardy et al. (Hardy et al., 2018), suggesting they are recurrent regulatory points in a conserved primary layer of starvation response.

### Mitochondrial bioenergetics and coordinated mito–nuclear signatures

Mitochondrial pathways emerged as key targets of starvation-induced differentiation, with nuclear-encoded mitochondrial genes being significantly overrepresented and involved in translation, central carbon metabolism, oxidative phosphorylation, lipid utilization, and mitochondrial maintenance. Differentiation at *Pdk*, a regulator of pyruvate entry into mitochondrial metabolism (Jeoung, 2015; Sugden and Holness, 2003), aligns with selection on metabolic routing during nutrient scarcity, while signals at *ND-ACP*, a core component of complex I, suggest adjustments in electron transport and redox balance (Toshniwal et al., 2019). Collectively, these loci form an interconnected bioenergetic network that controls metabolic efficiency, substrate flexibility, and mitochondrial integrity. Many show large allele-frequency shifts and strongly negative Tajima’s D, indicating sustained directional selection acting on linked genomic regions. The consistent differentiation within the mitochondrial genome, especially near the origin of replication (mtORI), further supports coordinated nuclear-mitochondrial responses. Additionally, predicted differences in mtORI structure and mtDNA copy number likely reflect selection on replication-related regions, supporting mito–nuclear coadaptation during prolonged starvation.

### Comparative signals in human orthologs of starvation-responsive fly genes

Cross-species analyses place these findings in a broader evolutionary context by asking whether starvation-responsive loci identified in flies correspond to differentiated orthologs in human populations included in our comparative PBS analysis. The increased survival under starvation and lipid storage observed in our starvation-selected fly lines are broadly compatible with the Thrifty Gene Hypothesis, which proposes that environments with high variance in caloric availability favored alleles that improve energy storage and metabolic efficiency (Neel, 1962). However, direct empirical support for this hypothesis in humans remains limited, and its genomic basis remains debated. In particular, the relative paucity of high-frequency, shared selective sweeps in human genomic datasets argues against a model in which starvation-related adaptation is driven by fixation of a few large-effect loci (Hancock et al., 2010; Speakman, 2008). Our fly results instead support a mixed genomic architecture, with recurrent involvement of some regulatory loci but a broader response distributed across multiple loci and pathways.

Using PBS analysis from the 1000 Genomes Project, we found significant enrichment of highly differentiated variants in human orthologs of starvation-responsive fly genes. This pattern is more consistent with coordinated allele-frequency shifts across functionally related genes than with repeated hard sweeps at the same loci. A smaller subset of regulatory loci was repeatedly implicated across analytical frameworks. In *Drosophila*, *S6k*, *sws*, rugose (*rg*), and *raskol* were recovered among drift-exceeding SNPs, low-heterozygosity intervals, and sweep-associated regions. Their human orthologs *RPS6KB*1/2, *PNPLA6*, *LRBA*, and *DAB2IP*, respectively, were likewise enriched for highly differentiated variants in the human comparison. The recurrence of these loci across replicated fly populations and in the ortholog-based human analysis is therefore consistent with repeated involvement of shared regulatory pathways under sustained nutrient limitation, rather than with strict locus-for-locus conservation of adaptation.

Notably, starvation-related variation at fly *S6k* is mostly noncoding, indicating selection on regulatory regions, and it is associated with decreased predicted RNA structural stability, implying possible post-transcriptional effects. Given the conserved role of reduced TOR–S6k signaling in promoting survival and autophagy during nutrient shortage (Kapahi et al., 2004), regulatory variation at this locus, i.e., less stable RNA-fold structure (Supplementary Figure S8), could also likely contribute to the increased longevity observed in this study and in previous starvation-evolution experiments (Bubliy and Loeschcke, 2005; Rion and Kawecki, 2007). However, testing these ideas experimentally is beyond the scope of this study. Additionally, *sws* encodes a phospholipase essential for membrane lipid homeostasis and neuronal maintenance in flies, which aligns with selection on membrane remodeling under prolonged starvation (Kmoch et al., 2015). *Rg*, a regulator of vesicle trafficking and autophagy, offers a plausible pathway by which nutrient stress may influence autophagic capacity (Shamloula et al., 2002), while Raskol integrates nutrient-sensitive insulin–Ras inputs with growth regulation through Ras–Rho signaling (Colgren and Nichols, 2019). In humans, these functional pathways are mirrored by *PNPLA6*, a fasting-induced lysophospholipase linked to basal mTORC1 output; LRBA, a regulator of autophagic flux; and *DAB2IP*, a RasGAP scaffold that reduces Ras signaling.

Our fly–human comparison is better understood as a comparative enrichment analysis rather than direct evidence of similar starvation adaptation. PBS captures branch-specific population differences but not the actual selective agent; differentiation at orthologous loci may be affected by diet, climate, pathogens, or demographic factors. The focal populations show increased branch-specific differentiation at orthologs of starvation-responsive fly genes, suggesting potential targets of evolutionary change even when nutrient stress is not the primary driver. However, the nonrandom enrichment of highly differentiated human variants in some orthologs of starvation-responsive fly genes indicates recurring involvement of metabolic and regulatory systems, such as insulin/TOR signaling, mitochondrial function, and lipid metabolism, while remaining cautious about assuming identical selective pressures across species.

In summary, our results suggest that adaptation to long-term starvation stress in *D. melanogaster* involves a complex genomic architecture that includes both strong regional signals of linked selection and widespread allele-frequency changes that go beyond neutral expectations. This pattern aligns with the idea that prolonged starvation affects multiple physiological systems rather than changes at a few isolated genetic loci. Repeatedly implicated genes and pathways include those involved in nutrient sensing, lipid metabolism, autophagy, and mitochondrial function, indicating that adaptation consistently targets fundamental processes related to energy balance and cellular maintenance. The partial overlap between starvation-responsive fly genes and their human orthologs supports the idea of pathway-level convergence under ongoing nutritional stress, even when gene-level overlap is limited. Overall, these findings offer an experimentally grounded framework for understanding how persistent nutrient stress can influence genomes through the combined effects of linked selection and polygenic adaptation.

## Material and Methods

### Fly collection and experimental evolution setup

Wild-type *Drosophila melanogaster* (n > 5,000) were collected during October–November 2021 from multiple locations across the Delhi National Capital Region (NCR), India (28.61° N, 77.23° E; altitude 216 m). To establish a genetically diverse base population, all wild-caught females were pooled and mass-bred for eight generations on standard yeast–cornmeal–agar medium under controlled laboratory conditions (25 °C, 70% relative humidity). Experimental evolution was initiated by allowing ∼1,000 gravid females to oviposit in four independent culture jars, from which one starvation-selected and one unselected control population were derived in a paired design, yielding four starvation-selected (SS_1–4_) and four control populations (C_1_–_4_). Starvation selection was imposed by starving ∼1,000 males and ∼1,000 females (3 days post-eclosion) per population to complete food deprivation on non-nutritive 2% agar until ∼75–80% mortality, after which the surviving individuals founded the next generation. Control populations were maintained in parallel on a standard diet without starvation stress. This selection regime was continued for 60 generations to establish stable starvation-selected lines. Prior to phenotypic and genomic analyses, all populations were maintained on standard food for six generations to minimize parental and environmental carryover effects.

### DNA extraction and whole-genome sequencing

Genomic DNA was extracted from pooled samples of 100 adult females per population (3 days post-eclosion) collected after 60 generations of experimental evolution, with four biological replicates per regime, using phenol:chloroform extraction followed by DNase-free RNase treatment. DNA quality and concentration were assessed prior to library preparation. Paired-end sequencing libraries (mean insert size ∼250 bp) were prepared using the Kapa HyperPlus kit and sequenced on an Illumina NovaSeq X Plus platform.

Raw reads were aligned to the *D. melanogaster* reference genome (FlyBase r6.06) using BWA (bwa mem, version 0.7.17) (Li and Durbin, 2009). Unmapped reads were removed, and BAM files were coordinate-sorted using SAMtools (v1.21). Duplicate reads were removed, and deduplicated BAM files were merged using Picard (v2.13.1) to generate final analysis-ready alignments for each population replicate. Variant calling was performed using GATK HaplotypeCaller (McKenna et al., 2010) with a pooled ploidy of 200 (--sample-ploidy 200, 100 diploid females). To retain a stringent, high-confidence variant set, variants near indels (within 5 nucleotides), clustered SNPs (3 SNPs within a window of 10 nucleotides), and other low-confidence sites were excluded. We additionally removed SNPs with low variant quality (QUAL < 30), evidence of strand bias (FS > 60), low sequencing depth (FORMAT/DP < 30), or low minor allele frequency per replicate (MAF < 0.05), ensuring a robust final call set for downstream analyses. After filtering, 1,508,941 high-confidence biallelic SNPs were retained.

To analyze mitochondrial variation, mitochondrial variants were subset from the genome-wide call sets by extracting variants overlapping mitochondrial coordinates using BEDTools (Quinlan and Hall, 2010; Ye et al., 2014). This yielded 70 mitochondrial SNPs, and pooled allele frequencies were estimated for each population to quantify mitochondrial allele-frequency variation. Finally, variants from both the nuclear and mitochondrial genomes were functionally annotated using ANNOVAR (Wang et al., 2010).

### Population-genetic statistics

Per-site heterozygosity was calculated as 2pq using population-level allele frequencies. Genome-wide heterozygosity was summarized in 100-kb sliding windows with a 2-kb step size to capture fine-scale variation in diversity. Windows with heterozygosity < 0.05 were classified as low-heterozygosity windows, and contiguous windows were merged to define extended regions of reduced genetic diversity.

Deviation from neutrality was assessed using Tajima’s D calculated in 10-kb windows with PoPoolation (Kofler et al., 2011). Nucleotide diversity (π) was calculated using custom R scripts. Selective sweep signatures were identified using Pool-HMM (Boitard et al., 2013), a hidden Markov model–based method tailored for Pool-Seq data, with parameters “-n 100 –c 10 –C 200 –q 20 –a unknown –k 0.001 –p”, using a pre-estimated site-frequency spectrum (specified via “-s”) for each chromosome. Sweep–low-heterozygosity regions were defined as regions supported jointly by low-heterozygosity patterns and Pool-HMM sweep signals.

### Drift-aware detection of selection

Allele frequencies were estimated for four control (C_1–4_) and four starvation-selected (SS_1–4_) populations using pooled whole-genome sequencing data, applying a diffusion-based approach to account for neutral drift, as previously described elsewhere (Hardy et al., 2018). This framework was used to distinguish allele-frequency shifts expected under neutral drift from those more consistent with selection in independently evolved starvation-selected populations. SNP-level allele counts were obtained following uniform quality filtering across all samples. Control allele frequencies were merged to compute mean control allele frequencies, which were converted to minor allele frequencies (MAF) and binned from 0.01 to 0.50 (rounded to the nearest hundredth). These control-derived allele frequencies were used as the starting frequency classes for modeling neutral divergence.

Effective population size (Ne) was inferred from allele-frequency variance among control populations, assuming divergence among controls reflects neutral drift. For each SNP, allele-frequency variance was computed across C replicates and then averaged within MAF bins; bins with MAF < 0.07 were excluded to reduce instability from rare alleles. Ne was then estimated using Kimura’s diffusion approximation (Kimura, 1955), separately for autosomes (Ne = 530) and the X chromosome (Ne = 461). Thus, chromosome-specific Ne estimates were used to account for differences in neutral drift dynamics between autosomes and the X chromosome.

Neutral allele-frequency evolution was modeled over 74 generations (8 laboratory-acclimation generations, 60 selection generations, and 6 common-garden generations) using diffusion-based Monte Carlo simulations. This interval corresponds to the total period over which control and starvation-selected populations were allowed to diverge prior to genomic comparison. For each starting MAF bin, drift trajectories were simulated from the corresponding initial MAF under the appropriate Ne, and the resulting distribution of absolute allele-frequency change (|ΔAF|) expected under drift alone was obtained. Observed allele-frequency shifts in each SS replicate were calculated relative to the mean allele frequency of the four control populations, and these values were evaluated against the corresponding simulated null distributions. The 99.999th percentile of simulated |ΔAF| distributions were used to define chromosome– and frequency-specific drift thresholds (a highly conservative genome-wide cutoff for identifying drift-inconsistent SNPs in Pool-seq data). For each starvation-selected replicate, observed |ΔAF| relative to the mean of the four control replicates was compared to the appropriate drift threshold, and SNPs exceeding threshold were classified as candidates under selection. Replicate-specific candidate sets were identified independently for each SS population. Finally, reproducibility across replicates was used to prioritize robust signals, with candidates consistently detected across multiple starvation-selected populations considered the strongest selection candidates. In particular, the intersection of candidate SNPs shared across all four starvation-selected replicates was used as the most conservative set for downstream functional analyses.

### Permutation-based significance testing

We used custom permutation tests in R to determine whether overlap among drift-filtered candidate loci and the representation of mito-nuclear genes among candidate genes were greater than expected under random sampling. For the drift-based analysis, SNP sets matching the observed candidate set sizes were repeatedly sampled without replacement from the filtered genome-wide SNP background, and the number of loci shared across sets was recorded for each iteration. For the mito-nuclear analysis, gene sets matched to the observed candidate set with respect to chromosome assignment and gene length were similarly sampled and evaluated for overlap with the MitoCarta reference list. Null distributions were generated from 100,000 permutations in each analysis, and empirical *P* values were calculated as the fraction of permutations equal to or exceeding the observed value.

### mtDNA content analysis

DNA was extracted from a 3-day-old female using a phenol–chloroform–isoamyl alcohol extraction and qPCR was performed on a Thermo Fisher QuantStudio 5 system with the G-Biosciences 2× SYBR Green qPCR mix.

### RNA secondary-structure prediction

To assess variant effects on RNA folding, reference and variant RNA sequences were analyzed with RNAfold (ViennaRNA Package) (Gruber et al., 2008) under default settings, and resulting minimum free energy (MFE; ΔG) values were compared. Specifically, the reference sequence was obtained from the *D. melanogaster* genome, and the variant sequence was generated by introducing population-specific SNPs (i.e., variants enriched in the starvation-selected populations). Reference and SNP-containing sequences were folded independently under identical conditions, without imposing structural constraints. For each sequence, RNAfold returned the predicted MFE secondary structure and ΔG; variant effects were evaluated by comparing ΔG as well as qualitative differences in the predicted base-pairing patterns and overall folding stability.

### Phenotypic and physiological characterization

For phenotypic and physiological characterization, 10 vials were generated from each population, and 10 individuals from each vial were used in the assays, yielding 100 individuals per population.

(i) *Starvation resistance assay* Starvation resistance was assayed using 3-day-old adult females from control and starvation-selected populations. Flies were briefly anesthetized, sorted into groups of 10 females per vial, and transferred to vials containing 2% non-nutritive agar to prevent desiccation while imposing complete nutritional deprivation. Vials were maintained at 25 °C under a 12 h light/12 h dark photoperiod, and agar vials were replaced every two days without anesthesia to minimize handling stress. Mortality was recorded at 4-h intervals until all individuals had died, and starvation resistance was quantified as survival time under food deprivation.
(ii) *Lifespan assay* Lifespan was measured using newly eclosed (1–4 h old) virgin female flies. Ten flies per replicate were maintained on standard food under laboratory conditions, with food replaced every 3 days. Mortality was recorded daily.
(iii) *Triacylglycerol (TAG) quantification* TAG levels were quantified using the CheKine™ Micro Triglyceride Assay Kit according to the manufacturer’s protocol. Ten adult flies per vial were homogenized on ice, clarified by centrifugation, and absorbance was measured at 420 nm. TAG content was normalized to total protein levels.

### Human population genomic analysis and Population Branch Statistic (PBS)

We tested whether genes under selection in starvation-evolved *D. melanogaster* also show elevated differentiation in human populations used as ancestry-linked proxies for broader regional nutritional histories. Population labels and autosomal genotype data were derived from Phase 3 of the 1000 Genomes Project (1000 Genomes Project Consortium 2015) (Auton et al., 2015). We analyzed four focal populations: Bengali from Bangladesh (BEB), Luhya in Webuye, Kenya (LWK), Sri Lankan Tamil in the UK (STU), and Yoruba in Ibadan, Nigeria (YRI); within population branch statistic (PBS) triplets that included a geographically or ancestrally proximate reference population and an outgroup. Specifically, BEB was analyzed with Gujarati Indians in Houston (GIH) as the reference and Finnish in Finland (FIN) as the outgroup; STU with Indian Telugu in the UK (ITU) and FIN; YRI with Esan in Nigeria (ESN) and Utah residents with Northern and Western European ancestry (CEU); and LWK with YRI and CEU (Alkorta-Aranburu et al., 2012). In each PBS triplet, the focal population was the lineage of interest, the reference population provided a closely related comparison, and the outgroup was used to polarize branch-specific allele-frequency divergence. These populations were treated as ancestry-linked proxies for broader regional nutritional histories rather than as evidence of famine exposure in the sequenced individuals. This interpretation is strongest for BEB, given the well-documented famine history of Bengal/Bangladesh and broader evidence linking chronic energy stress in South Asia to metabolic constraint (De Waal, 2018; Famine histories, 2024; Wells et al., 2016). The African comparison populations were likewise justified at the regional level by recurrent famine associated with drought, poverty, conflict, displacement, and political instability (Bush, 1996; De Waal, 2018; Macrae and Zwi, 1992).

Fly candidate genes were defined using two complementary lines of evidence: genes in sweep–low-heterozygosity regions, and genes linked to drift-filtered candidate SNPs. We mapped these sets to human orthologs using DIOPT (Garcia and Korhonen, 2020), retaining only high-confidence assignments (DIOPT score >7) with strong and method-consistent support. When a fly gene mapped to multiple high-confidence human orthologs, all were retained. This yielded two human ortholog sets corresponding to the two fly-derived candidate classes.

For each focal–reference–outgroup triplet, we calculated per-SNP Weir–Cockerham *F*_ST_ and converted these values to the Population Branch Statistic (PBS) as described elsewhere (Alkorta-Aranburu et al., 2012; Jha et al., 2016), thereby estimating lineage-specific allele-frequency divergence in the focal population relative to the reference and outgroup. For each comparison, differentiated loci were defined empirically as SNPs in the upper 5% and 1% tails of the genome-wide PBS distribution, respectively. PBS-tail SNPs were assigned to genes based on overlap with annotated human genes, and for each focal population, we identified those that fell within the two human ortholog sets derived from fly candidate genes. We then tested whether SNPs in these orthologous genes were enriched in the upper 5% and upper 1% PBS tails relative to the genome-wide autosomal background for that population, using two-sided binomial tests. To evaluate repeatability, we quantified the number of orthologous genes containing at least one high-PBS SNP in each population and measured overlap among populations and between the two fly-derived ortholog classes. Genes recurrently recovered across populations and candidate classes were prioritized as the strongest candidates for conserved starvation-associated differentiation across species.

## Data availability

The sequencing data generated in this study have been submitted to NCBI under project ID PRJNA1422946. All other data supporting the findings of this study are available within the article and its Supplementary Material.

## Acknowledgments

We sincerely thank Professor Gyaneshwer Chaubey (Banaras Hindu University, India) for insightful discussions and thoughtful comments on the manuscript. This work is supported by the Department of Biotechnology (DBT), Government of India, under grant no. BT/PR44574/MED/30/2401/2022 (Chronic Disease). GY acknowledges fellowship support from the University of Delhi for the doctoral program. We also acknowledge the publicly available human genomic resources generated by the 1000 Genomes Project Consortium, which provided an important comparative framework for this study.

## Supplementary Information details

### Supplementary Tables

**Table S1.** Sequencing quality and mapping summary for control and starvation-selected populations.

**Table S2.** Genomic distribution and density of identified variant sites across functional annotation categories.

**Table S3.** Chromosome-wise summaries of heterozygosity and Tajima’s D in control and starvation-selected populations.

**Table S4.** Numbers of selective sweep regions identified on each chromosome in individual control and starvation-selected replicate populations.

**Table S5.** Chromosome-wise counts of selective sweep regions in control and starvation-selected populations.

**Table S6.** Overlap between selective sweep regions and low-heterozygosity blocks across the genome.

**Table S7.** Candidate genes located within sweep–low-heterozygosity regions.

**Table S8.** Gene-wise sweep scores in control and starvation-selected populations.

**Table S9.** GO enrichment results for genes in sweep–low-heterozygosity regions.

**Table S10.** List of candidate genes post filtering genetic drift.

**Table S11.** Go terms of genes filtered post genetic drift using Gowinda.

**Table S12.** List of genes overlapped between sweep–low-heterozygosity regions and drift filtered genes.

**Table S13.** Candidate genes overlapping with the MitoCarta mitochondrial gene set and their GO terms.

**Table S14.** Primer list

**Table S15.** Functional annotations of genes shared among the top 5% PBS outliers across all analyzed human populations.

**Table S16.** Comparison of the present findings with related studies.

### Supplementary Figures

**Figure S1.** Genome-wide reductions in heterozygosity in starvation-selected populations.

**Figure S2.** Starvation selection increases heterozygosity variance and generates large, recurrent selective sweep regions.

**Figure S3.** Genome-wide reduction in nucleotide diversity in starvation-selected populations.

**Figure S4.** Heterozygosity and Tajima’s D profiles across the major chromosome arms aligned with selective-sweep signatures.

**Figure S5.** Functional enrichment of genes under starvation-driven selection.

**Figure S6.** Frequency-dependent estimates of effective population size and neutral drift dynamics.

**Figure S7.** Starvation-associated variants alter predicted RNA secondary structure of mtORI.

**Figure S8.** Starvation-associated variants alter predicted RNA secondary structure of *S6k*.

## Supplementary figures

**Supplementary Figure S1.**
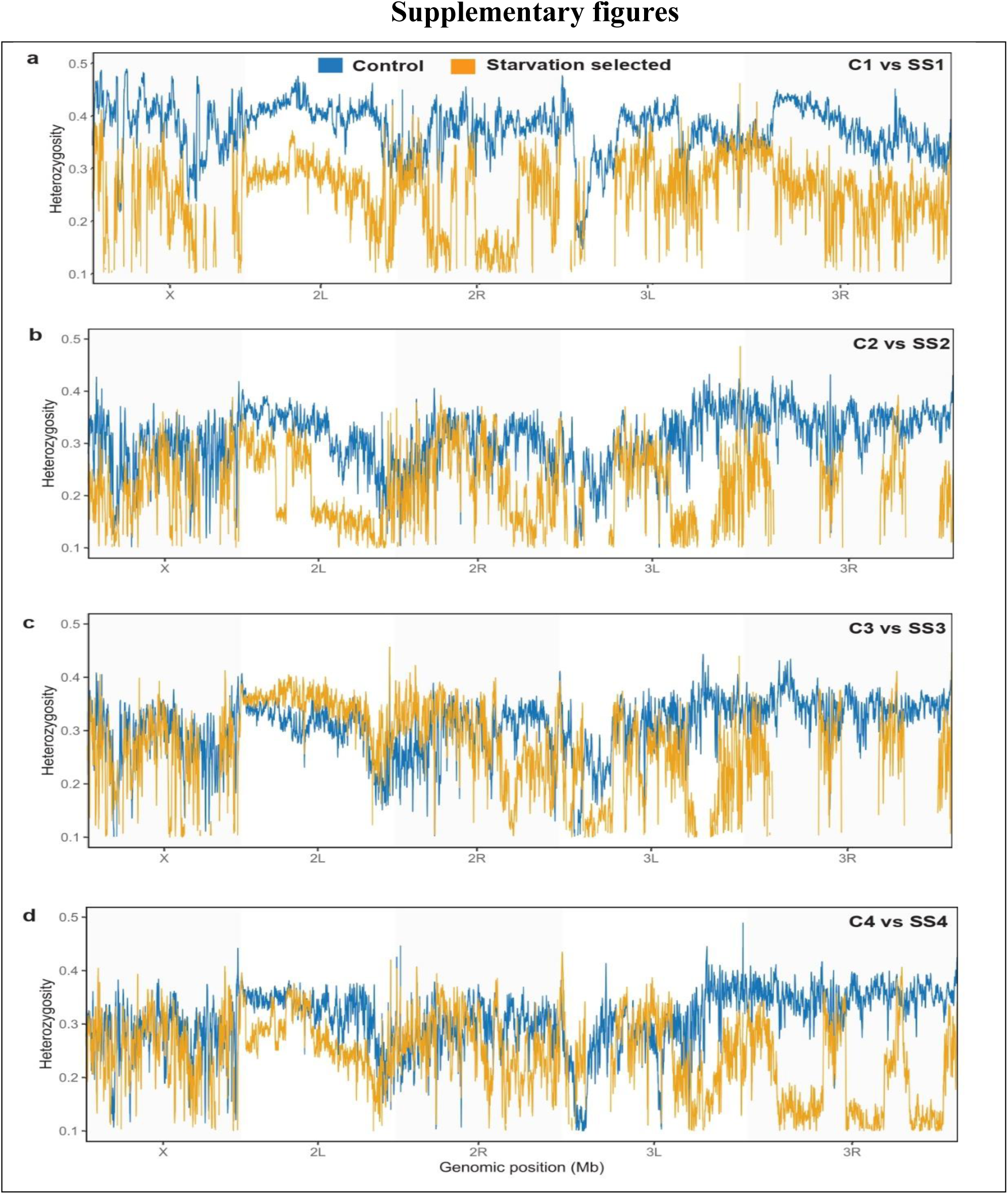
Genome-wide reductions in heterozygosity in starvation-selected populations. (a–d) Genome-wide heterozygosity profiles estimated using sliding windows (100-kb windows with a 2-kb step size) for each control (C) and starvation-selected population (SS) pair (C1–SS1 through C4–SS4). SS_1-4_ populations (orange) exhibit extensive, spatially coherent reductions in heterozygosity across multiple chromosomal regions, consistent with strong directional selection. In contrast, C_1-4_ populations (blue) maintain relatively uniform heterozygosity across the genome. The x-axis denotes genomic position along chromosome arms X, 2L, 2R, 3L, and 3R.

**Supplementary Figure S2.**
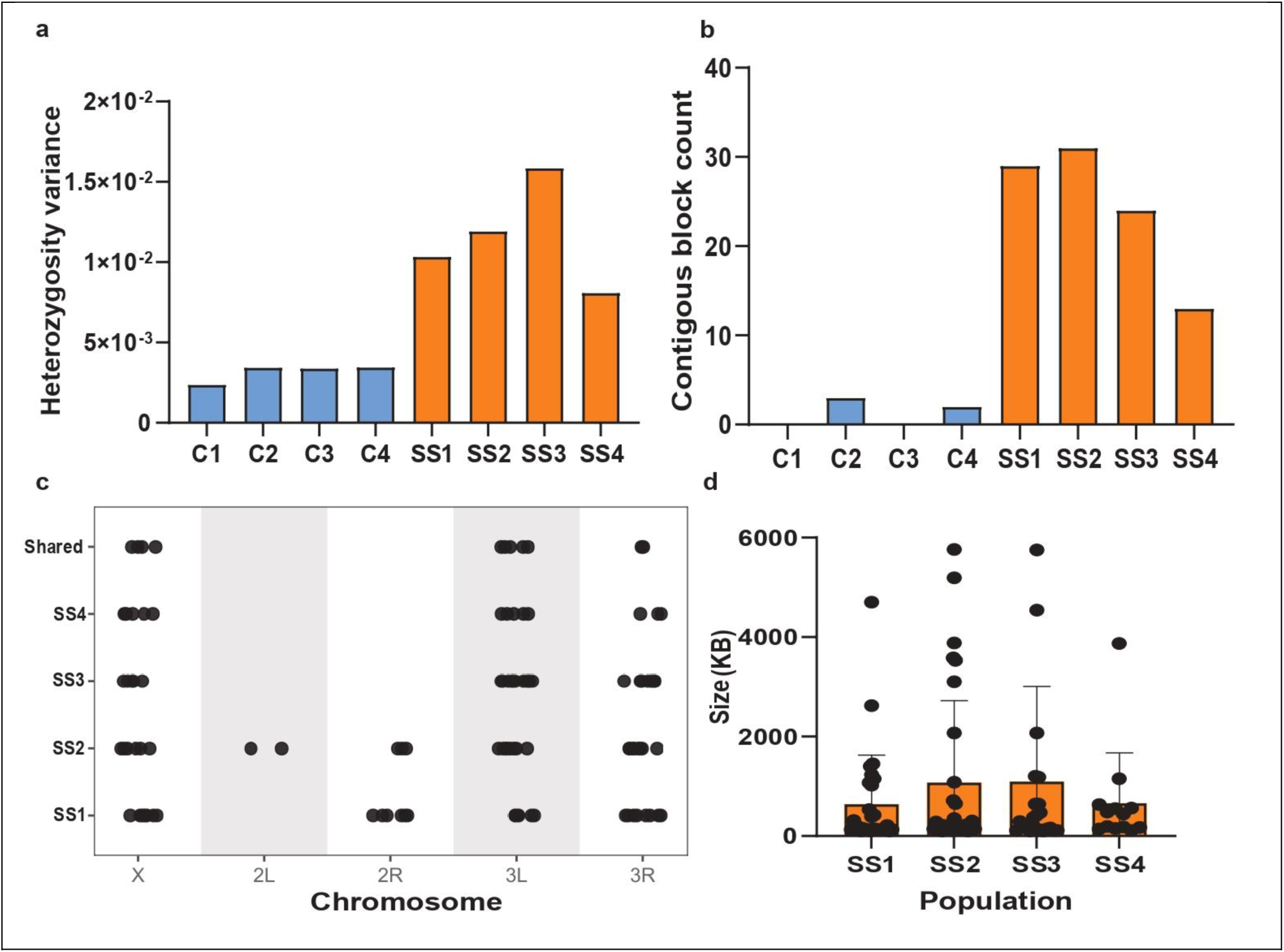
Starvation selection increases heterozygosity variance and generates large, recurrent selective sweep regions. (a) Starvation-selected (SS) populations show a pronounced increase in heterozygosity variance relative to controls (σ²_SS = 0.008–0.015 vs. σ²_Control = 0.002–0.003; Levene’s test, *P* < 1 × 10⁻⁶). (b) Number of contiguous low-heterozygosity blocks (H < 0.05) identified per population. Starvation-selected populations harbour dramatically more such blocks (734–3,027 per line) than controls populations. (c) Chromosomal distribution of low-heterozygosity blocks across SS replicates, showing multiple regions shared among independent populations. Shared blocks are indicated in the top row. (d) Size distribution of contiguous low-heterozygosity blocks in SS populations. Sweep sizes range from 0.6 to 1.1 Mb ± SEM and are significantly larger than those observed in control populations.

**Supplementary Figure S3.**
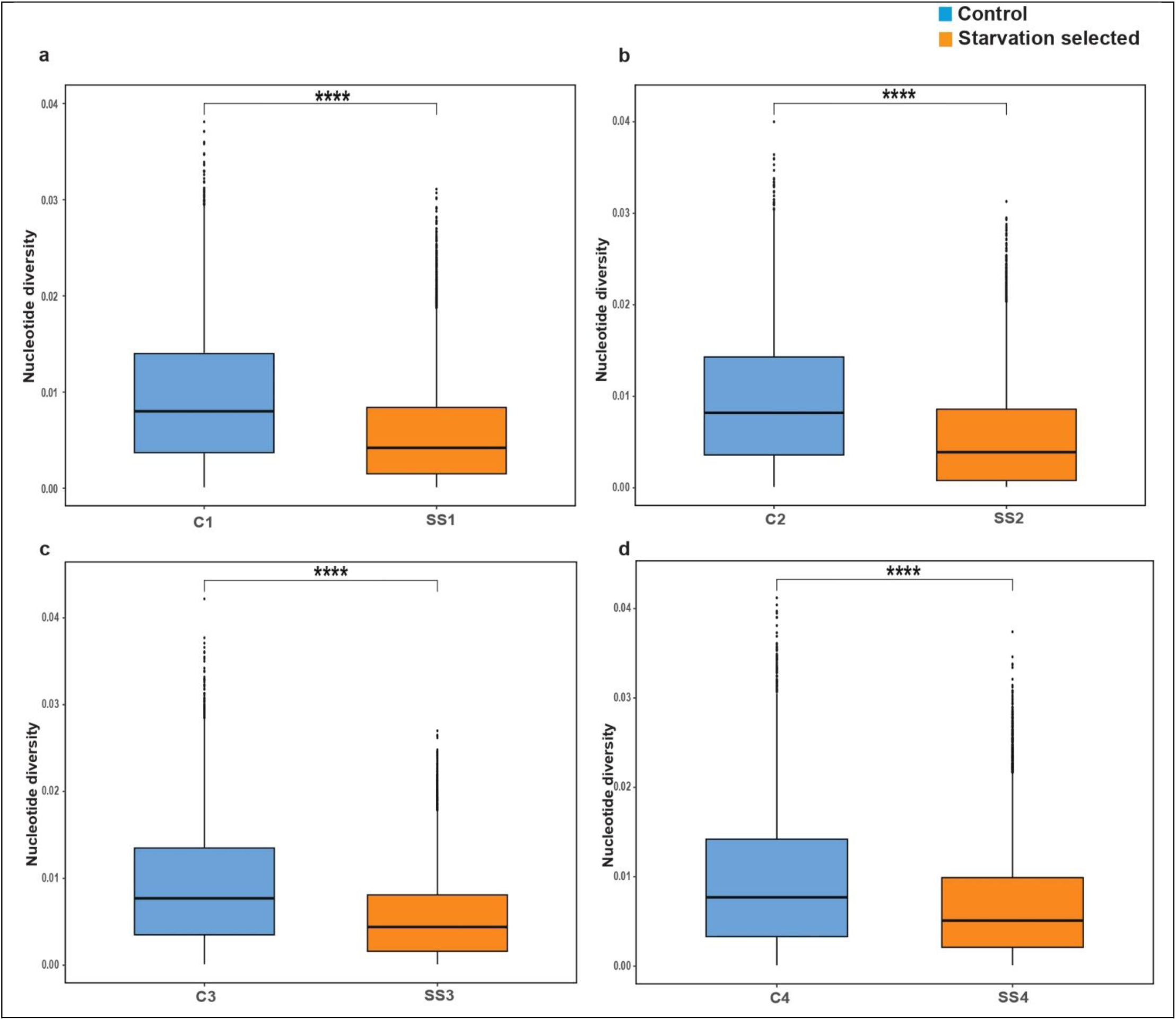
Genome-wide reduction in nucleotide diversity in starvation-selected populations. (a–d) Genome-wide nucleotide diversity (π) estimated for each matched control–starvation-selected (SS) population pair (C_1_–SS_1_ through C_4_–SS_4_). In all four comparisons, SS populations exhibit significantly lower nucleotide diversity than their corresponding C populations, consistent with widespread loss of genetic variation under sustained directional selection. Boxplots show median, interquartile range, and whiskers extending to 1.5× the interquartile range; points represent individual windows. Statistical significance was assessed using the Wilcoxon rank-sum test; \*\*\**P* < 0.0001.

**Supplementary Figure S4.**
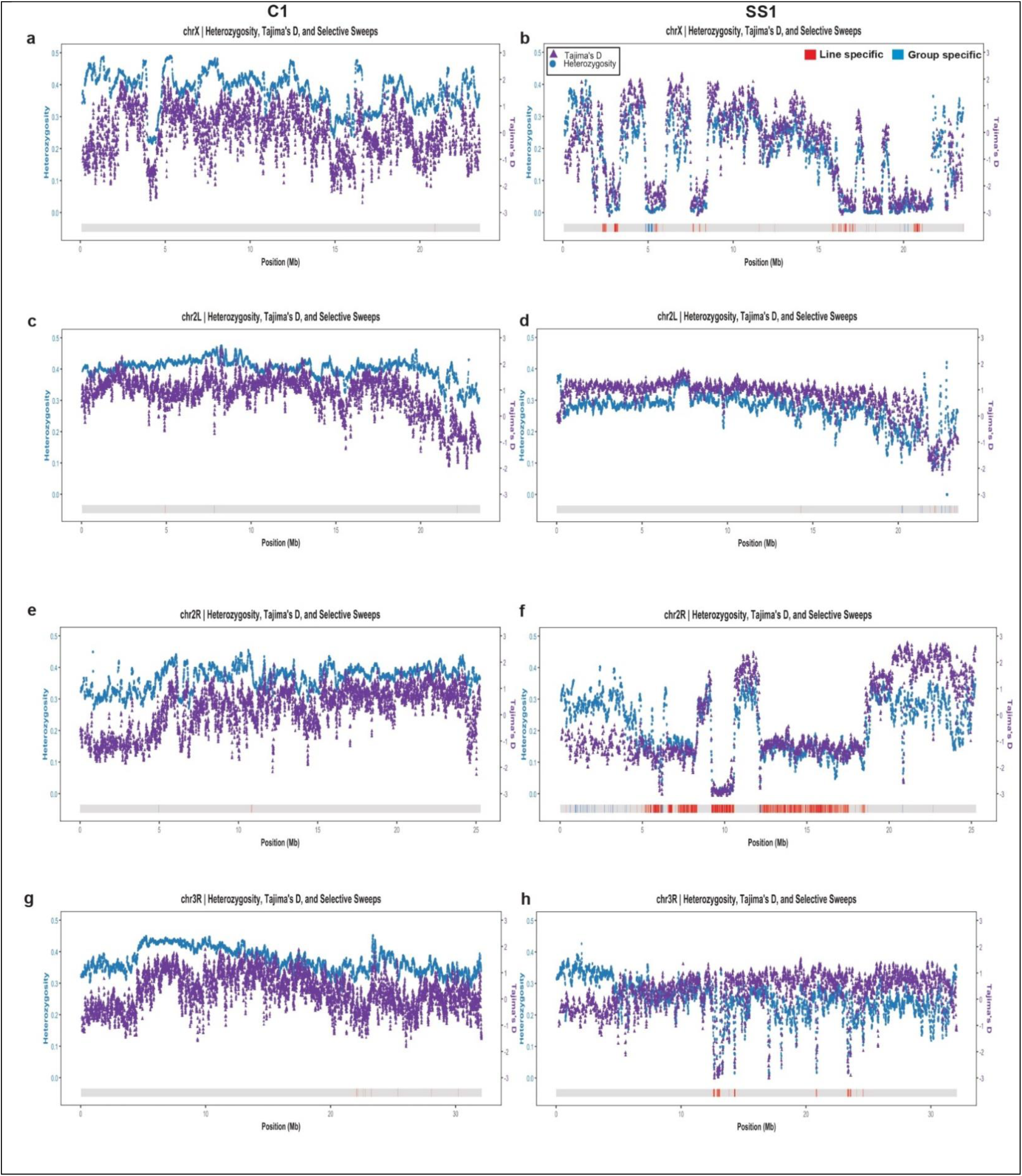
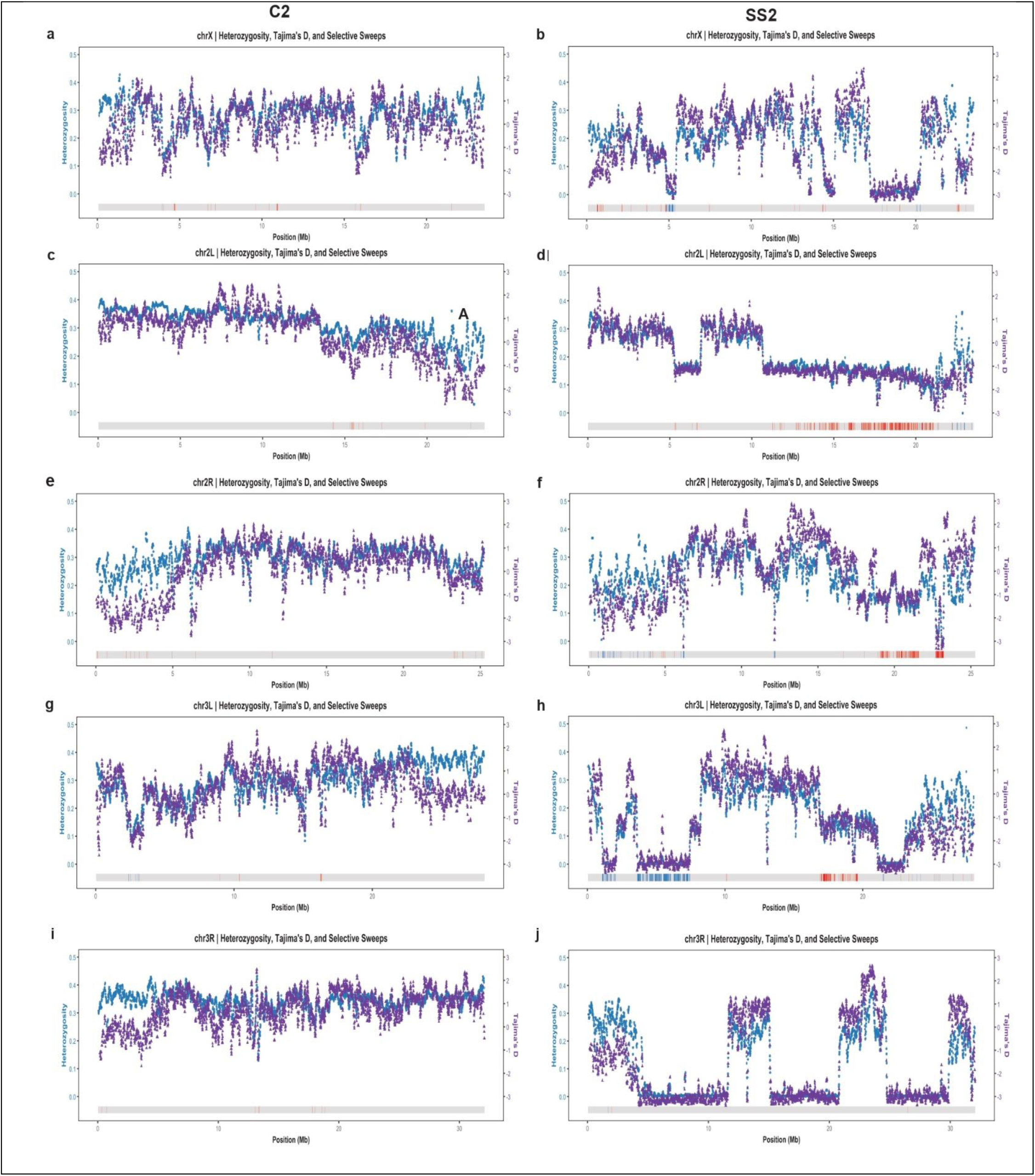
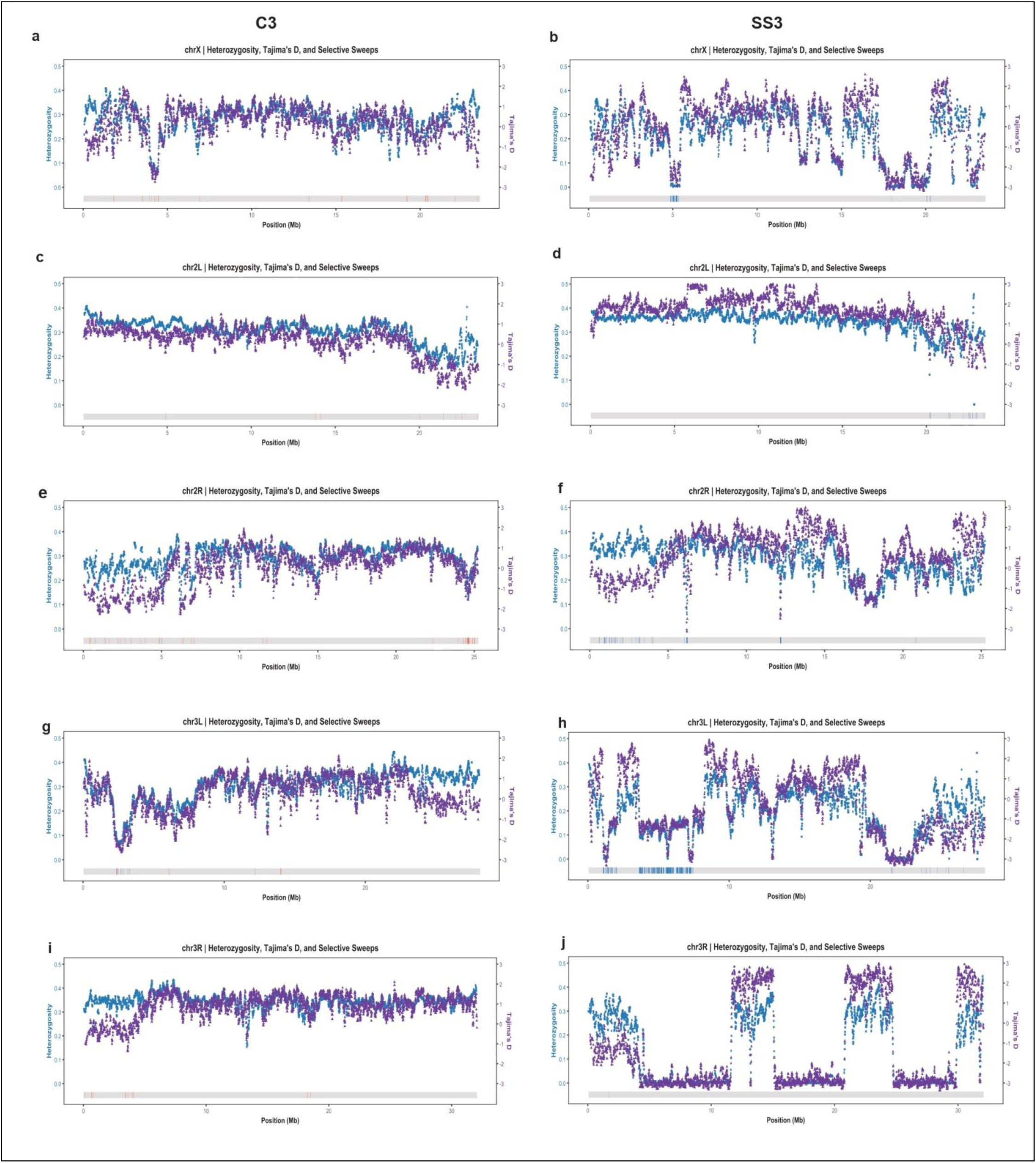
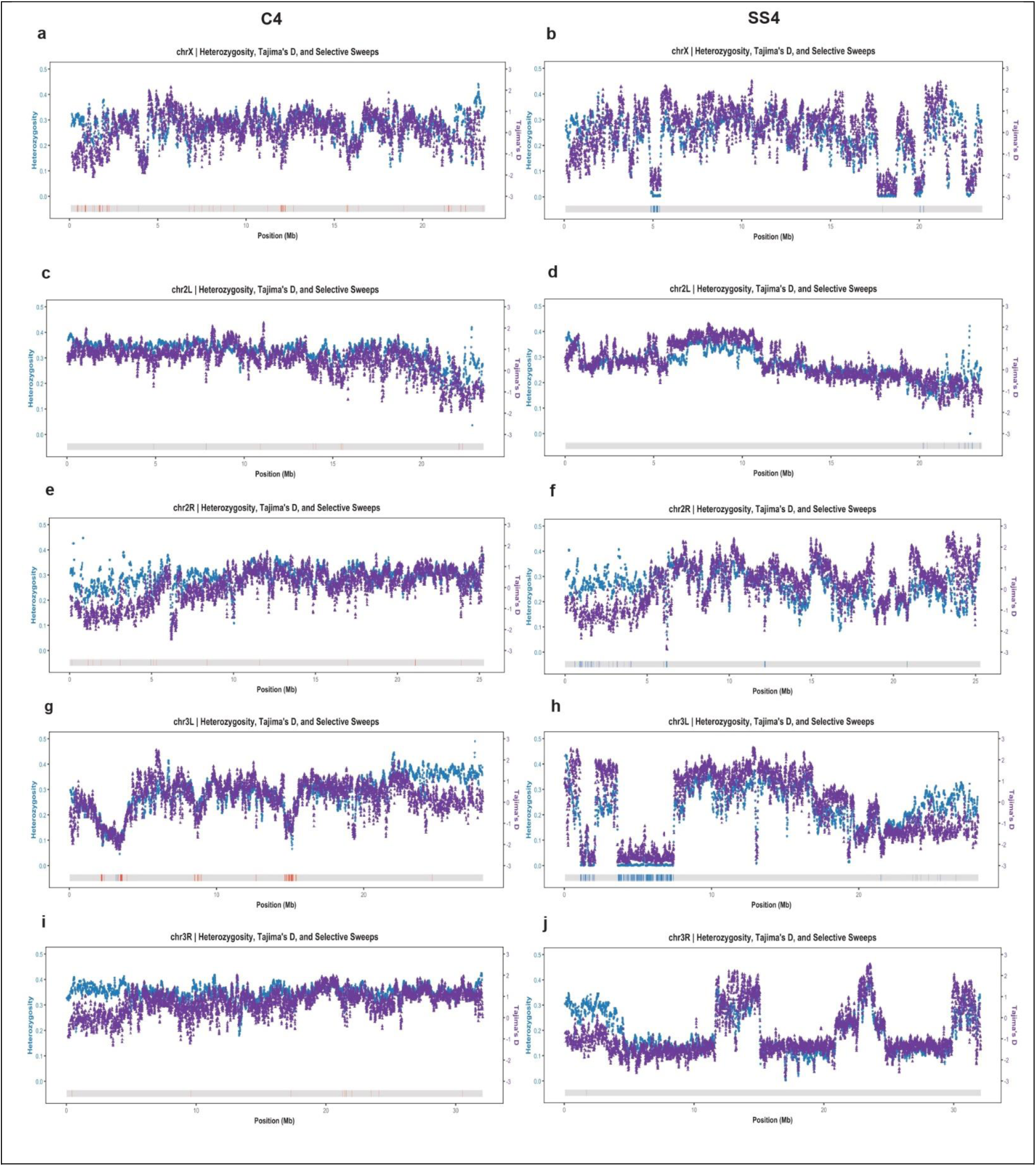
Heterozygosity and Tajima’s D profiles across the major chromosome arms aligned with selective-sweep signatures. Genome-wide estimates of heterozygosity (blue points) and Tajima’s D (purple triangles) are shown for control and starvation-selected (SS) populations. Colored blocks mark inferred sweep intervals: sweeps shared across all SS populations (blue), line-specific sweeps detected in individual SS populations (red), and regions with no sweep signal (gray).

**Supplementary Figure S5.**
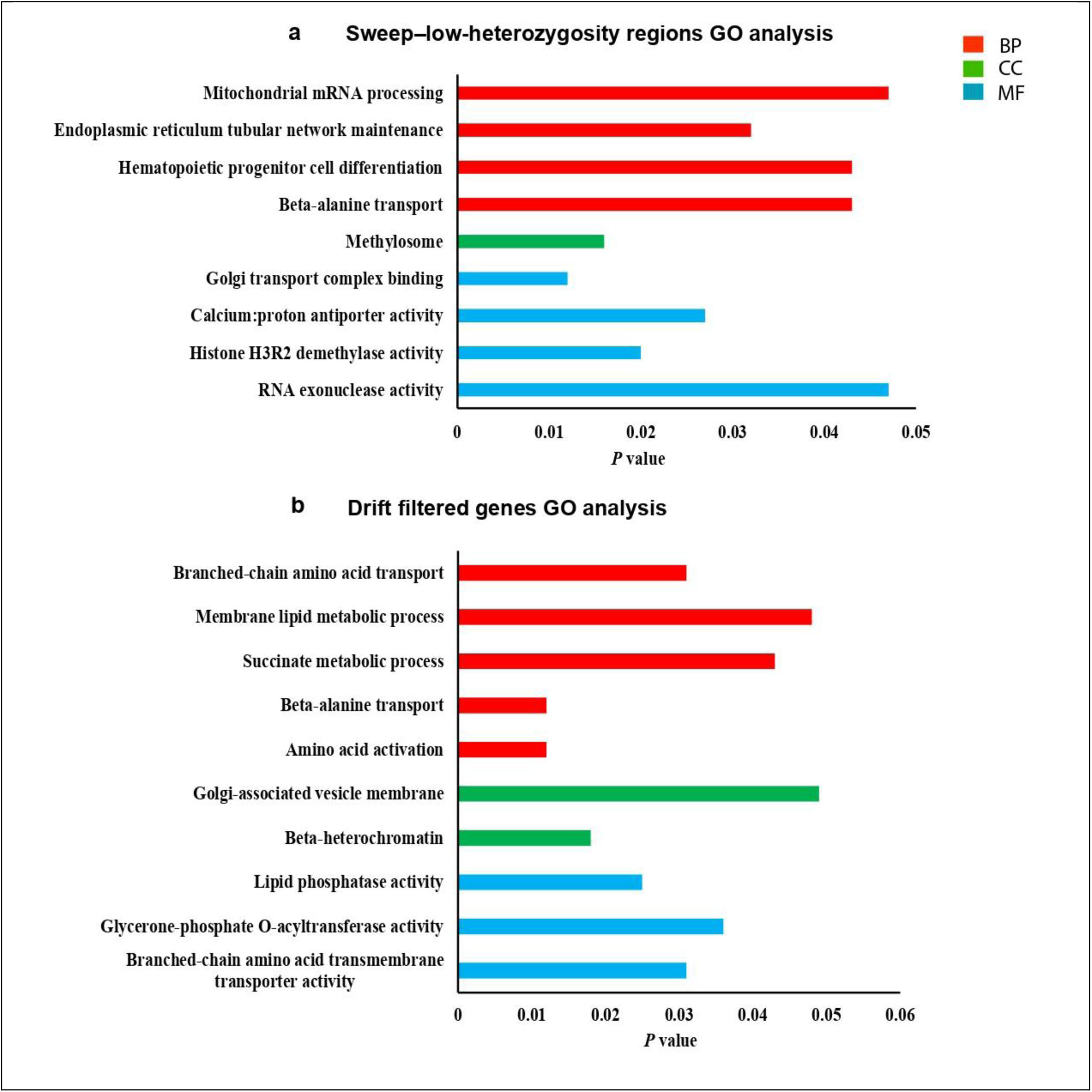
Functional enrichment of genes under starvation-driven selection. (a) GO enrichment analysis of genes within sweep–low-heterozygosity regions reveals distinct functional signatures, including pathways involved in RNA processing, chromatin modification, organellar maintenance, intracellular transport, and mitochondrial mRNA processing. (b) Gene Ontology (GO) enrichment analysis of drift-filtered genes identifies significant overrepresentation of biological process (BP), molecular function (MF), and cellular component (CC) categories related to amino acid transport and activation, lipid metabolic processes, mitochondrial function, and membrane-associated compartments. Bars indicate *P* values for significantly enriched GO terms.

**Supplementary Figure S6.**
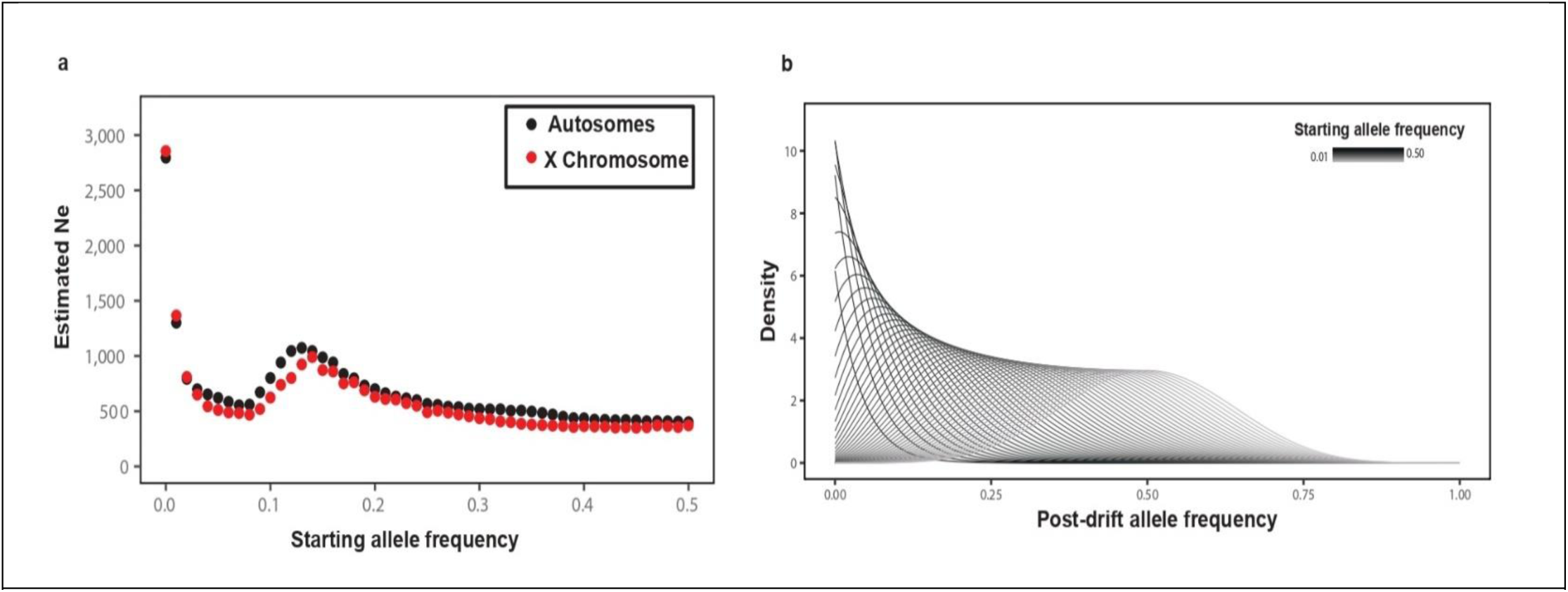
Frequency-dependent estimates of effective population size and neutral drift dynamics. (a) Relationship between starting allele frequency and the effective population size (Nₑ) estimated from control populations based on variance in allele frequencies across replicates. Autosomal sites (black) and X-linked sites (red) show similar frequency-dependent patterns, with Ne peaking at low initial allele frequencies (∼0.05) and declining at higher frequencies, consistent with reduced sampling variance at intermediate allele frequencies. (b) Posterior distributions of allele-frequency change under neutrality generated by Monte Carlo simulations for discrete starting-frequency bins. Darker curves correspond to higher initial allele frequencies. The pronounced skew toward low post-drift frequencies for rare alleles illustrates the asymmetric nature of drift-driven dynamics in pooled population sequencing.

**Supplementary Figure S7.**
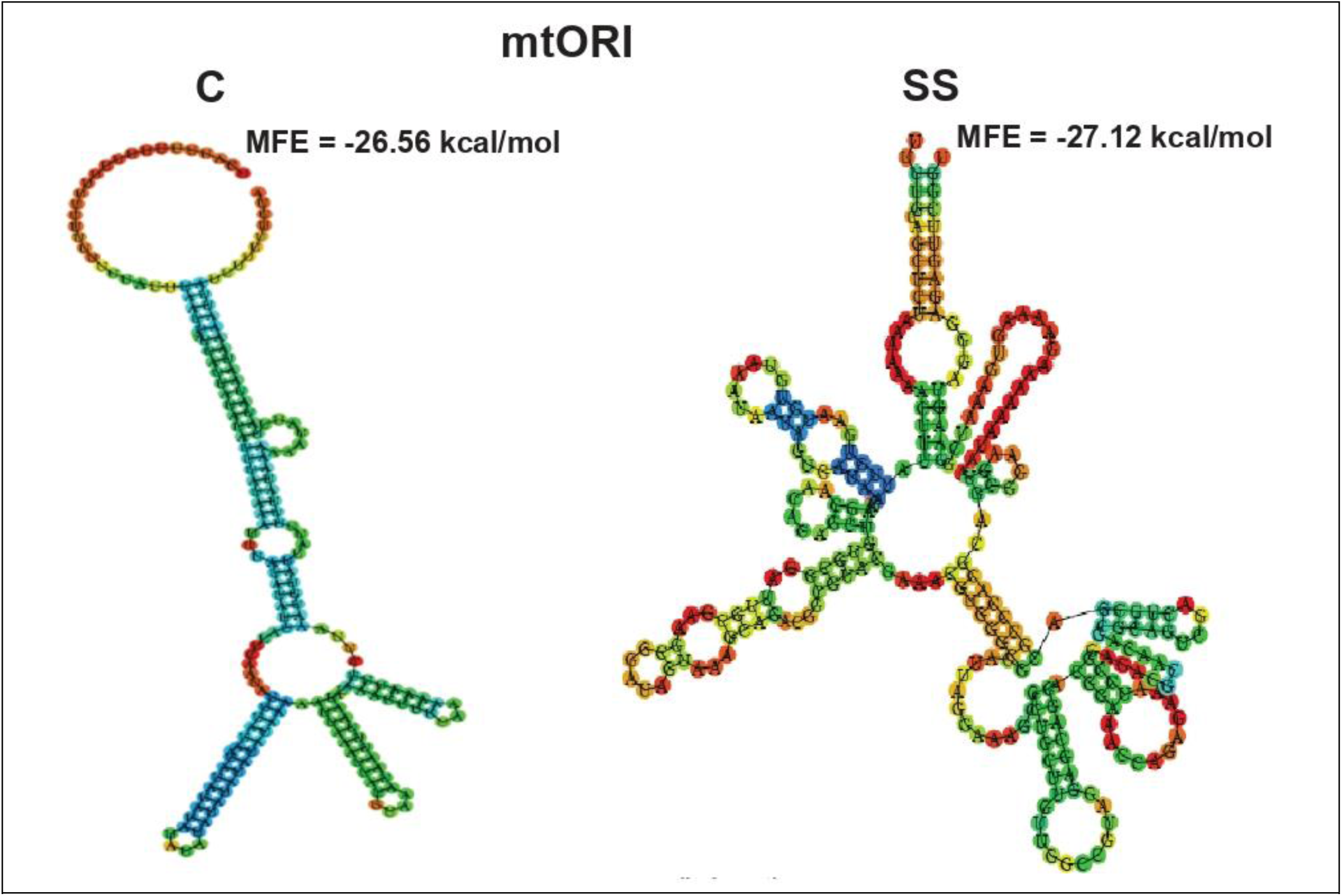
Starvation-associated variants alter predicted RNA secondary structure of mtORI. Predicted mtORI RNA structures for control (C; left) and starvation-selected (SS; right) populations were generated using identical folding parameters. Variants alter folding topology and local stem–loop architecture, with MFE values shown (C: −26.56; SS: −27.12 kcal/mol). Base pairing is color-coded by structural context (red–orange, highly paired; yellow–green, intermediate).

**Supplementary Figure S8.**
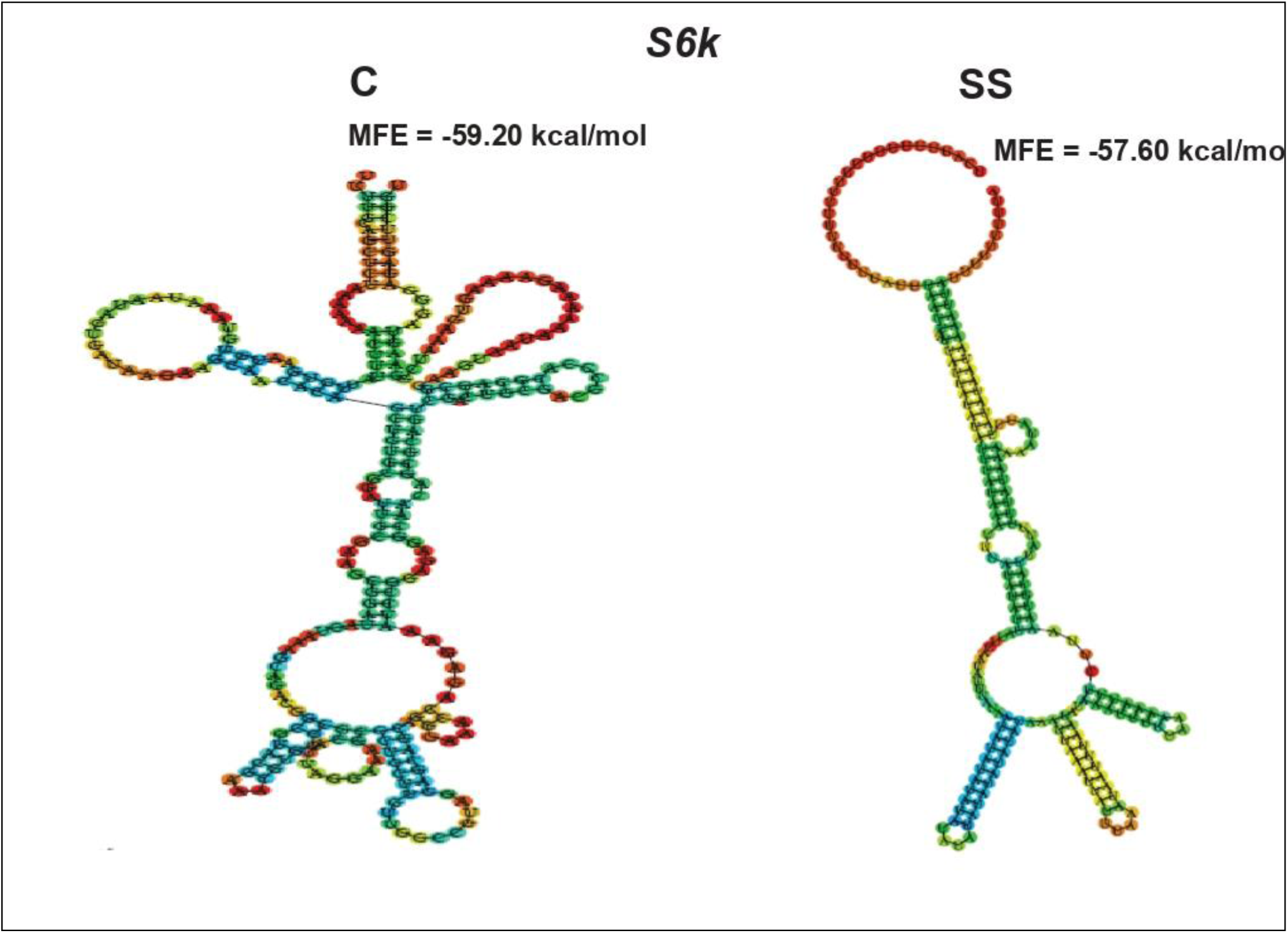
Starvation-associated variants alter predicted RNA secondary structure of *S6k*. Predicted RNA secondary structures for *S6k* in control (C; left) and starvation-selected (SS; right) populations. Structures were generated from reference (control) and starvation-associated variant DNA sequences using identical folding parameters. Starvation-associated variants produce alterations in RNA folding topology and local stem–loop architecture, suggesting potential effects on RNA stability, processing, or translation. Base pairing is color-coded by structural context, with red–orange indicating highly stable, strongly paired regions and yellow–green indicating regions of intermediate stability.

